# A high-throughput all-optical platform for mapping gene perturbations to electrophysiology

**DOI:** 10.64898/2026.11.09.765233

**Authors:** Jinfan Zhang, Daozheng Gong, Alec M. Barrios, Daniel G. Itkis, Keemaya Sahni, Hannah Fandl, Takeshi Uenaka, Russell T. Walton, Sarah E. Plutkis, Luke D. Lavis, Paul C. Blainey, Marius Wernig, Ralda Nehme, Richard T. Lee, Ernest Fraenkel, Samouil L. Farhi, Adam E. Cohen

**Affiliations:** Department of Chemistry and Chemical Biology, Harvard University, Cambridge, MA; Platform Innovation Labs, Broad Institute of MIT and Harvard, Cambridge, MA; Stanley Center for Psychiatric Research, Broad Institute of MIT and Harvard, Cambridge, MA; Broad Institute of MIT and Harvard, Cambridge, MA; Department of Biology, Massachusetts Institute of Technology, Cambridge, MA; Department of Biological Engineering, Massachusetts Institute of Technology, Cambridge, MA; Department of Stem Cell and Regenerative Biology, Harvard University, Cambridge, MA; Institute for Stem Cell Biology C Regenerative Medicine, Stanford University School of Medicine, Stanford, CA; Department of Pathology, Stanford University School of Medicine, Stanford, CA; Department of Physics, Harvard University, Cambridge, MA; Janelia Research Campus, Howard Hughes Medical Institute, Ashburn, VA; Koch Institute for Integrative Cancer Research at MIT, Cambridge, MA

**Author notes:** Equal contribution (Jinfan Zhang, Daozheng Gong, Alec M. Barrios).

## Abstract

Electrical phenotyping remains a bottleneck in linking genes to cellular function. We developed a high-throughput all-optical electrophysiology platform combining an engineered human induced pluripotent stem cell (hiPSC) line, wide-field voltage imaging, and automated analysis. The hiPSC line integrates inducible NGN2, CRISPR interference, and degron-gated expression of Voltron2 and CheRiff, enabling neuronal differentiation, targeted gene knockdown, and optical electrophysiology. The microscope records approximately 1,500 spiking glutamatergic neurons at 800 Hz across a 4x4 mm field, with larger fields supporting approximately 3,000 neurons at 660 Hz. Automated analysis extracts single-cell voltage traces and quantifies firing patterns, action-potential waveforms, and subthreshold dynamics. Arrayed knockdown of five sodium- and potassium-channel genes produced distinct electrophysiological phenotypes. The engineered hiPSC line also supported optical electrophysiology recordings from motor neurons, astrocytes, and cardiomyocytes. These results establish a scalable platform for high-throughput electrophysiological phenotyping and functional genomics in hiPSC-derived cells.

## Introduction

Electrical activity underlies communication, development, and homeostasis across many cell types^1,2^. In neurons, firing patterns and action-potential waveforms emerge from the coordinated activity and spatial arrangements of ion channels, transporters, receptors, and intracellular signaling pathways^3^. Mapping how genes shape these electrical phenotypes would help connect molecular mechanisms to neuronal function and disease.

Cellular excitability emerges from the collective activity of many interacting conductances, together with processes that control channel abundance, trafficking, and modulation.^4–6^ Homeostatic compensation, functional redundancy, and nonlinear interactions can make the consequences of a genetic perturbation difficult to predict^4,8,9^. Direct measurements of electrophysiological phenotypes are therefore needed to determine how individual genes contribute to cellular electrical function.

Existing electrophysiological methods impose a tradeoff between throughput and information content. Patch-clamp electrophysiology directly measures membrane potential or ionic currents, but scaling to large populations of neurons remains difficult. Calcium imaging and multielectrode arrays can monitor activity from many cells simultaneously, but do not directly report action-potential waveforms or subthreshold dynamics^10,11^. A scalable assay that combines controlled stimulation with cell-resolved voltage measurements would directly connect genetic perturbations to electrical phenotypes.

All-optical electrophysiology (Optopatch) combines an optogenetic actuator with a genetically encoded voltage indicator for optical stimulation and voltage recording^12^. Optopatch has been applied to measurements of neuronal excitability, pharmacological responses, and disease-associated electrophysiological phenotypes^13–15^. Early Optopatch constructs used the very dim near-infrared fluorescence of Archaerhodopsin-derived voltage indicators. Even with ultrawide-field optics, as in the Firefly microscope^16^, two factors limited the field of view (FOV): 1) the requirement for high-intensity laser illumination (> 100 W/cm^2^) restricted the FOV to ∼3 mm^2^. Above this area, laser-induced sample heating affected neural function. 2) at 16-bit digitization, only ∼200 rows of the camera chip could be read at 1 kHz. As a result, Optopatch recordings were limited to an FOV of 0.5 × 4 mm at 1 kHz^16^, or 0.8 × 4 mm at 500 Hz.^14^

Chemigenetic voltage indicators such as Voltron2 provide bright fluorescence and sub-millisecond voltage responses^17^. Labeling with the red-shifted HaloLigand dye JF_608_ enables spectrally independent voltage imaging and CheRiff stimulation.^18^ The increased brightness of Voltron2 permits voltage imaging with LED illumination at 2 – 5 W/cm^2^, making larger imaging fields practical. Usable area then depends on illumination uniformity, optical performance, camera readout, and signal extraction.

Combining Optopatch with genetic perturbations could enable mapping of genes to electrophysiology at high throughput. Calcium-based CRISPR screens have identified genes that regulate neuronal activity.^19^ Combining CRISPR perturbations with Optopatch recordings could reveal effects on waveform and subthreshold dynamics as well as firing. Several technical challenges must be overcome: 1) stable and homogeneous Optopatch expression throughout differentiation; 2) sufficient light collection, image quality, and camera throughput across a large field; and 3) automated analysis that separates overlapping cells and converts large movies into quantitative phenotypes. Thus, the cellular substrate, optical instrumentation, and analysis pipeline must be engineered together to optimize overall measurement throughput and information content.

Here, we address these three challenges. We engineered a hiPSC line that combines inducible neuronal differentiation, CRISPR interference (CRISPRi)-mediated gene perturbation, and stable integration of Voltron2 and CheRiff with degron-gated control. An ultrawide-field microscope records approximately 1,500 active neurons across a 4 × 4 mm field at 800 Hz, or approximately 3,000 neurons across a 5.1 × 5.1 mm field at 660 Hz. An automated analysis pipeline identifies active cells, extracts single-cell voltage traces, detects action potentials, and quantifies electrophysiological features.

An arrayed benchmark targeting five sodium- and potassium-channel genes revealed distinct effects on evoked firing, action-potential waveform, adaptation, and post-stimulation repolarization. The same engineered hiPSC line also supported Optopatch measurements in differentiated motor neurons, astrocytes, and cardiomyocytes. Together, these components establish a scalable platform for functional genomics of electrophysiology in hiPSC-derived cells.

## Results

### A stable hiPSC line enables Optopatch and CRISPRi

A genetic screen of neuronal electrophysiology requires reproducible differentiation, efficient and sustained genetic perturbation, and uniform expression of both the voltage reporter and optogenetic actuator. We therefore sought to generate a hiPSC line in which all three capabilities were genomically encoded and maintained throughout neuronal differentiation. We started from a widely used and characterized hiPSC line, WTC11 carrying doxycycline-inducible NGN2 at the AAVS1 safe-harbor locus for rapid and homogeneous differentiation into excitatory neurons, together with CAG-driven dCas9-BFP-KRAB at the CLYBL locus for CRISPRi (Coriell GM29371; Fig. 1a) ^7^.

**Figure 1.**
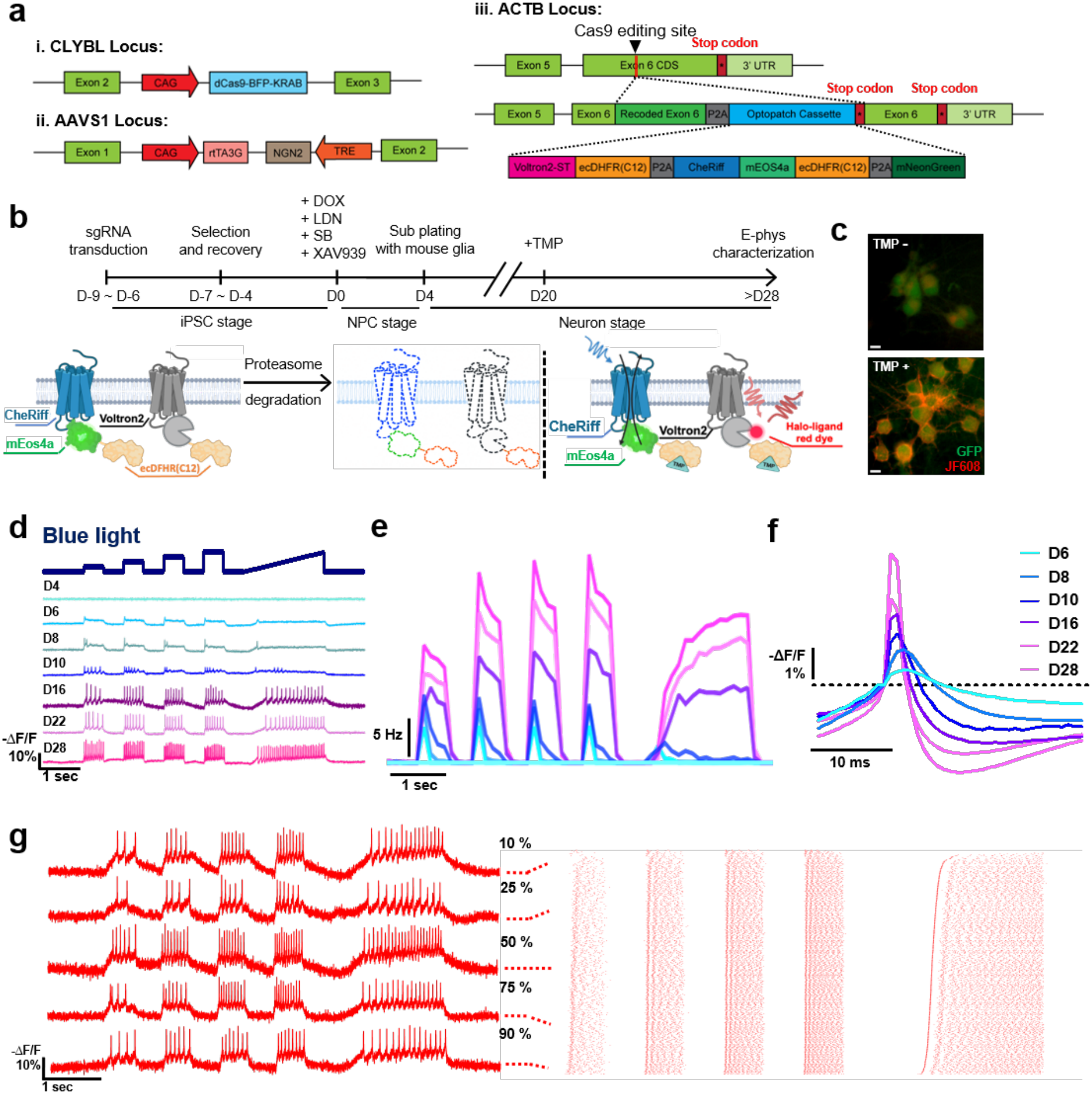
A multifunctional engineered human iPSC line enables optical electrophysiology of iPSC-derived neurons. **(a)** Genomic configuration of the parental CRISPRi-NGN2 iPSC line and the ACTB-targeted Optopatch cassette. The parental line contained (i) dCas9-BFP-KRAB at the CLYBL locus, (ii) doxycycline-inducible NGN2 at the AAVS1 locus. We added (iii) degron-tagged Voltron2 and CheRiff at the ACTB locus. Trimethoprim (TMP) stabilized the ecDHFR(C12)-tagged Optopatch proteins. **(b)** Experimental timeline for targeted gene knockdown and optical electrophysiology measurements. **(c)** Representative images showing TMP-dependent Optopatch expression in DIV28 neurons following TMP addition at DIV20. Scale bar 10 µm. **(d)** Representative Optopatch recordings during neuronal differentiation. **(e)** Population-average firing-rates for the same stimulation protocol as in (d). D4 NPCs, *N* = 35; D6 neurons, *N* = 163; D8, *N* = 193; D10, *N* = 53; D16, *N* = 82; D22, *N* = 117; D28, *N* = 171. **(f)** Spike-triggered average spike waveforms across neuronal differentiation. **(g)** Representative voltage traces from neurons at the indicated percentiles of excitability. Right: raster plots of all imaged neurons during Optopatch protocol in (d), sorted by optical rheobase. N = 1,046 neurons.

The Optopatch components comprised soma-targeted variants of the chemigenetic voltage indicator Voltron2 and the blue-light-activated channelrhodopsin CheRiff^12,17,20^. Optopatch constructs have commonly been delivered by viral vectors, which leads to several-fold cell-to-cell variation in reporter and actuator expression. This heterogeneity stresses the dynamic range of the imaging system, and leads to variable photocurrents across cells exposed to the same illumination intensity. Initial efforts to maintain stable Optopatch expression in iPSCs led to impaired cell survival over extended passaging, consistent with reports that sustained expression of membrane-bound rhodopsins can be poorly tolerated in pluripotent stem cells^21^. In addition, exogenous expression cassettes can undergo transcriptional silencing during neuronal differentiation, preventing stable transgene expression^22,23^.

We tested three strategies for achieving strong and sustained Optopatch expression after differentiation, while minimizing expression burden in proliferating iPSCs. First, we used TALEN-mediated insertion of an hSyn-driven Optopatch cassette, flanked by UCOE and cHS4 anti-silencing elements, at the available CLYBL locus. Second, we used PiggyBac transposition to introduce multiple copies of a similarly protected hSyn-driven cassette^23^ (Fig. S1a). In both strategies, the neuron-selective hSyn promoter was used to restrict Optopatch expression to differentiated neurons and thereby minimize expression in iPSCs. Third, we used SLEEK-mediated insertion of Optopatch into the endogenous ACTB locus, coupling expression to the constitutive β-actin transcript through a self-cleaving P2A peptide^24^ (Fig. 1a). In this design, both Voltron2 and CheRiff were fused to ecDHFR(C12) degrons, enabling post-translational control of protein abundance by trimethoprim (TMP)^25^. This strategy maintained low Optopatch protein levels during iPSC expansion while permitting robust accumulation after TMP addition in differentiated cells. In all three strategies, CheRiff was fused to the green-to-red photoswitchable fluorescent protein mEOS4a^26^, providing a fluorescent marker of CheRiff expression and enabling UV based photo labeling and optical cell-selection applications^27^ (a feature not used in the present work). These constructs also contained mNeonGreen as a fluorescent marker of successful genomic integration and cassette expression, as Optopatch protein expression was intentionally suppressed at the iPSC stage.

NGN2-induced excitatory neurons were differentiated by doxycycline-mediated induction in the presence of SB431542, LDN193189, and XAV939 for 3 days, followed by replating onto astrocyte-containing imaging plates on Day 4 and subsequent maturation in neuronal medium supplemented with BDNF, CNTF, and GDNF^28^ (Fig. 1b; Methods). The CLYBL-targeted line expressed its fluorescent marker in iPSCs but lost most Optopatch expression after neuronal differentiation, with little Voltron2 or GFP fluorescence detectable by DIV28, consistent with silencing during differentiation. In contrast, both the polyclonal PiggyBac line and the monoclonal ACTB-SLEEK line retained Optopatch expression after differentiation. In the ACTB-SLEEK line, Optopatch fluorescence was minimal in the absence of TMP, indicating effective suppression by the ecDHFR(C12) degrons (Fig. 1c). Addition of 10 μM TMP beginning at DIV20 produced strong Optopatch expression by DIV28, as measured by JF_608_-labeled Voltron2 fluorescence (Fig. 1b–c; Fig. S2). Both the PiggyBac and ACTB-SLEEK lines were maintained for more than 20 passages without an apparent loss of viability or expression stability.

We next compared the uniformity of Optopatch expression and electrophysiological responses among ACTB-SLEEK neurons, PiggyBac neurons, and neurons transduced with Optopatch lentivirus after differentiation (Figs. S1 and S2). Flow cytometry showed that ACTB-SLEEK neurons had a more uniform distribution of Voltron2 and GFP fluorescence than either the PiggyBac or lentiviral populations (Fig. S1b-c, Table S1). We then compared functional responses under the same spatially homogeneous blue-light stimulation protocol. PiggyBac neurons showed lower signal-to-noise ratios and more variable evoked firing, whereas lentivirally transduced neurons produced stronger signals but retained substantial cell-to-cell heterogeneity (Table S2). ACTB-SLEEK neurons gave strong signals and had the lowest cell-to-cell variability in firing rates (Fig. 1g; Fig. S3a-c), with a narrower distribution of total evoked spike counts under a standard step-ramp blue light waveform (Fig. S3d, Table S2). Under ramped blue light, ACTB-SLEEK neurons showed a more consistent optical rheobase and a reproducible rise in firing rate with stimulus strength, whereas PiggyBac and lentiviral populations varied more in both threshold and stimulus-response relation (Fig. S3e,f, Table S2). Together, these measurements show that integration of Optopatch at the ACTB locus, combined with TMP-dependent gated expression, provides stable and high level Optopatch expression and homogeneous functional measurements.

Because Optopatch abundance in the ACTB-SLEEK line is controlled by TMP rather than by a differentiation-stage-specific promoter, expression can be induced at different stages of neuronal maturation. We used this feature to track electrophysiological maturation during differentiation. Immature neurons showed weak or limited optogenetically evoked responses, whereas progressively more mature neurons generated larger and more frequent action potentials during a standardized blue-light stimulation protocol (Fig. 1d,e; Fig. S4). Action-potential waveform also changed substantially with maturation: spike-triggered averages increased in amplitude and became sharper, and the afterhyperpolarization became progressively more pronounced (Fig. 1f). By DIV22–28, neurons exhibited robust stimulus-dependent firing and mature action-potential waveforms (Fig. 1g).

### A high-throughput imaging platform for Optopatch phenotyping of iPSC-derived neurons

To maximize the number of neurons recorded in each acquisition, we built an ultrawide-field microscope for high-speed, large-area Optopatch imaging (Fig. 2a). High-power blue and red LED illumination were combined with a dichroic mirror, spatially homogenized, and projected onto a digital micromirror device (DMD), enabling spatially patterned optogenetic stimulation and excitation. The microscope permitted independent and near-simultaneous patterning in each wavelength, by subdividing each camera frame into epochs with distinct DMD patterns, and synchronously switching the LED light sources to only illuminate the corresponding DMD patterns (here we only used wide-field illumination in both colors). A large dichroic mirror directed the combined illumination onto the sample. Fluorescence from Voltron2-JF_608_ was collected through the same objective, transmitted through the dichroic and an emission filter, and recorded on a high-speed sCMOS camera. A low-magnification objective combined with a long-focal-length tube lens provided an effective magnification of 2.3×.

**Figure 2.**
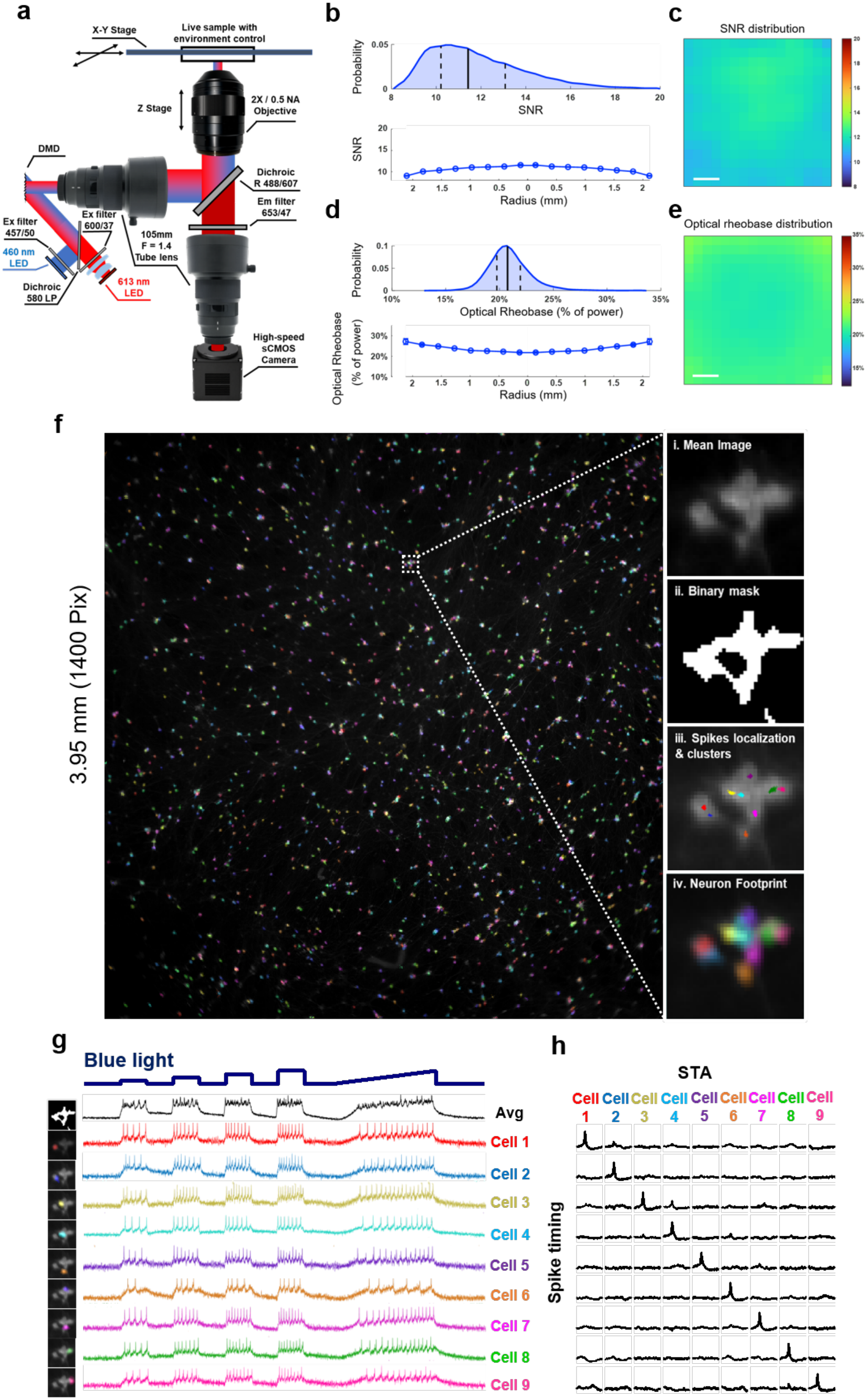
Ultrawide-field optical electrophysiology enables high-throughput recording and automated extraction of single-neuron voltage traces. **(a)** Schematic of the ultrawide-field Optopatch microscope. Blue light for CheRiff stimulation and red light for Voltron2 excitation are combined and patterned with a single digital micromirror device (DMD), directed to the sample through a low-magnification objective, and Voltron2 fluorescence is collected onto a high-speed sCMOS camera. **(b)** Top: Distribution of single- spike signal-to-noise ratio (SNR) across cells. Solid and dashed vertical lines indicate median and interquartile range, respectively. Bottom: mean SNR across the imaging field. Error bars, SEM. **(c)** Map of mean spike SNR across the field of view. Scale bar 0.5 mm. **(d)** Top: Distribution of optical rheobase across cells. Optical rheobase is expressed as a percentage of maximum blue-light power. Solid and dashed vertical lines indicate median and interquartile range, respectively. Bottom: mean optical rheobase across the imaging field. Error bars, SEM. **(e)** Map of optical rheobase across the field of view. Scale bar, 0.5 mm. **(f)** Automated identification of active neurons using Activity Localization Imaging (ALI). The representative 1,400 × 1,400-pixel field contains 1586 activity-defined neuronal footprints, with individual neurons shown in different pseudocolors. Insets illustrate the analysis workflow for a representative region: (i) mean fluorescence image; (ii) binary mask defining a contiguous region containing one or more cells; (iii) localization and clustering of detected spikes; and (iv) activity-defined neuronal footprints obtained by averaging spike-associated -ΔF images within each cluster. **(g)** Representative extraction of voltage traces from spatially overlapping neurons. The black trace shows the spatially averaged signal from the selected region; colored traces show the individually extracted voltage signals. Footprints shown at left. **(h)** Quantification of signal separation among the nine neurons shown in (g). Each row shows spike-triggered averages (STAs) calculated using the spike times of one neuron and applied to the traces of all nine neurons.

We performed a quantitative analysis of tradeoffs in camera bit-depth, dynamic range, and speed, and identified conditions which enabled a 6-fold increase in FOV with minimal increase in noise compared to prior efforts (Fig. S5). The key insight was that if total photon counts were kept high, one could truncate the least-significant bits with little noise penalty, because these bits were dominated by shot noise. Specifically, if the number of photon counts is *N*, the shot noise scales as *N*^1/2^. Consequently, if these counts are represented as an *m*-bit number (*m* = ceil[log_2_ *N*]), then the bottom *m*/2 bits are dominated by shot noise – discarding these bits costs little in SNR. Hence, we operated the camera (Teledyne Kinetix) in 8-bit acquisition mode, which increased the pixel readout rate approximately sixfold relative to 16-bit acquisition. To maintain high photon capacity, we used a custom deep-well firmware mode that supported signals of up to approximately 4,000 photoelectrons per pixel per frame. The reduced bit-depth only increased the noise by ∼40% relative to the 16-bit mode (Fig. S5b). This configuration enabled voltage imaging at 800 Hz over 1,400 × 1,400 pixels, corresponding to an approximately 4 × 4 mm FOV or 660 Hz over 1,800 × 1,800 pixels, corresponding to a 5.1 × 5.1 mm FOV (Fig. S5a).

The microscope was controlled using Luminos software^29^, which synchronized sample translation, autofocus, optical stimulation, image acquisition, and data saving. Including stage positioning and data transfer, the system acquired approximately 150 fields of view per hour. At a typical yield of approximately 1,500 spiking neurons per field, this corresponds to a measurement throughput of approximately 225,000 spiking neurons per hour.

We next quantified the SNR and spatial uniformity of Optopatch measurements across the field of view. A typical FOV (1,400 × 1,400 pixels) contained ∼1,500 spiking neurons or 3,000 spiking neurons at 2000 × 2500 pixels. Near the periphery of this larger imaging area, optical aberrations distorted neuronal footprints, but high-quality voltage traces could still be extracted (**Movies 1 G 2**). The population distribution of spike SNR was centered at 11.4 (IQR: 10.2 - 13.1, N = 9075 neurons). Spatial mapping showed a 23% systematic decrease in SNR at the edges of the FOV (Fig. 2b,c).

We then characterized the variability in optical excitability, quantified via optical rheobase, defined as the blue-light intensity at which a neuron first fired during a ramp stimulus. The population distribution of optical rheobase was centered at 20.8% of maximum blue power (IQR: 19.8-21.9%, N = 9075 neurons). Because CheRiff photocurrent depends on blue-light intensity, systematic spatial variation in rheobase provides a functional measure of stimulation uniformity. Optical rheobase increased by 24% near the edges of the FOV (Fig. 2d,e). Together, the weak spatial dependence of spike SNR and optical rheobase indicates that the microscope provides sufficiently uniform red-light excitation and fluorescence collection, as well as blue-light optogenetic stimulation, for population-scale comparisons across a multi-millimeter field of view.

The large field of view created an analysis challenge: each recording contained fluorescence signals from thousands of closely spaced and partially overlapping neurons. We therefore developed an automated analysis pipeline based on Activity Localization Imaging (ALI)^30^ to identify active cells and extract their individual voltage traces (Fig. 2f). Structural fluorescence images were first thresholded to identify contiguous regions containing one or more neurons. Within each region, the pixel-averaged fluorescence trace was used to detect candidate action potentials. Because Voltron2 fluorescence decreases upon depolarization, spike-associated images were represented as −ΔF so that action potentials appeared as positive signals. For each detected spike, we calculated the fluorescence-weighted spatial centroid, and spikes with nearby centroids were grouped into candidate neurons. Averaging the spike-associated −ΔF images within each cluster generated an activity-defined spatial footprint for each neuron. This procedure separated individual neurons even when neighboring cells overlapped extensively in the structural fluorescence image (Fig. 2f).

We then used these activity-defined footprints to recover each neuron’s fluorescence time course from the full movie. The movie was approximated as the product of a spatial matrix, describing the contribution of each neuronal footprint to each pixel, and a temporal matrix containing the corresponding fluorescence traces. With the spatial footprint matrix fixed, the temporal components were estimated by pseudoinverse, yielding single-neuron voltage traces throughout the recording (Fig. S6). In the representative 1,400 × 1,400-pixel field, the pipeline identified and extracted voltage traces from approximately 1,600 active neurons in a single 800-Hz acquisition. To test whether the unmixing procedure could resolve signals from spatially overlapping neurons, we examined a representative region containing nine activity-defined cells (Fig. 2g). The spatially averaged signal from this region contained a mixture of spikes from multiple neurons, whereas footprint-based extraction recovered distinct voltage traces with different spike timings from each cell. We further quantified signal separation by triggering the trace of every cell using the spike times detected in each of the nine neurons (Fig. 2h). Each neuron showed a strong spike-triggered response when triggered by its own spike times, whereas cross-triggered responses in neighboring cells were substantially smaller. Normalizing the cross-triggered SNR to the self-triggered SNR produced a diagonal matrix, demonstrating that the analysis pipeline effectively separated voltage signals from closely spaced and overlapping neurons.

Together, the wide-field optical system and automated activity-based analysis pipeline enabled high-speed electrophysiological measurements from thousands of neurons per acquisition while preserving spatially uniform stimulation, sufficient voltage-imaging SNR, and single-cell signal separation.

### CRISPRi perturbation of ion channels reveals gene-specific changes in neuronal excitability and action-potential dynamics

To test whether the platform could resolve electrophysiological phenotypes produced by genetic perturbations, we perturbed ion channels with established roles in neuronal excitability. sgRNAs were delivered by lentivirus using the CROP-seq-Multi-V2 vector, with two sgRNAs targeting each gene^31^ (Fig. S7a). We targeted the potassium-channel genes KCNT2 and KCNQ2 and the sodium-channel genes SCN2A, SCN8A, and SCN9A, together with a non-targeting control (NTC). Transcript measurements confirmed efficient knockdown of each target gene, with minimal effects on the expression of the other ion-channel genes examined (Fig. 3a; Fig. S7b).

**Figure 3.**
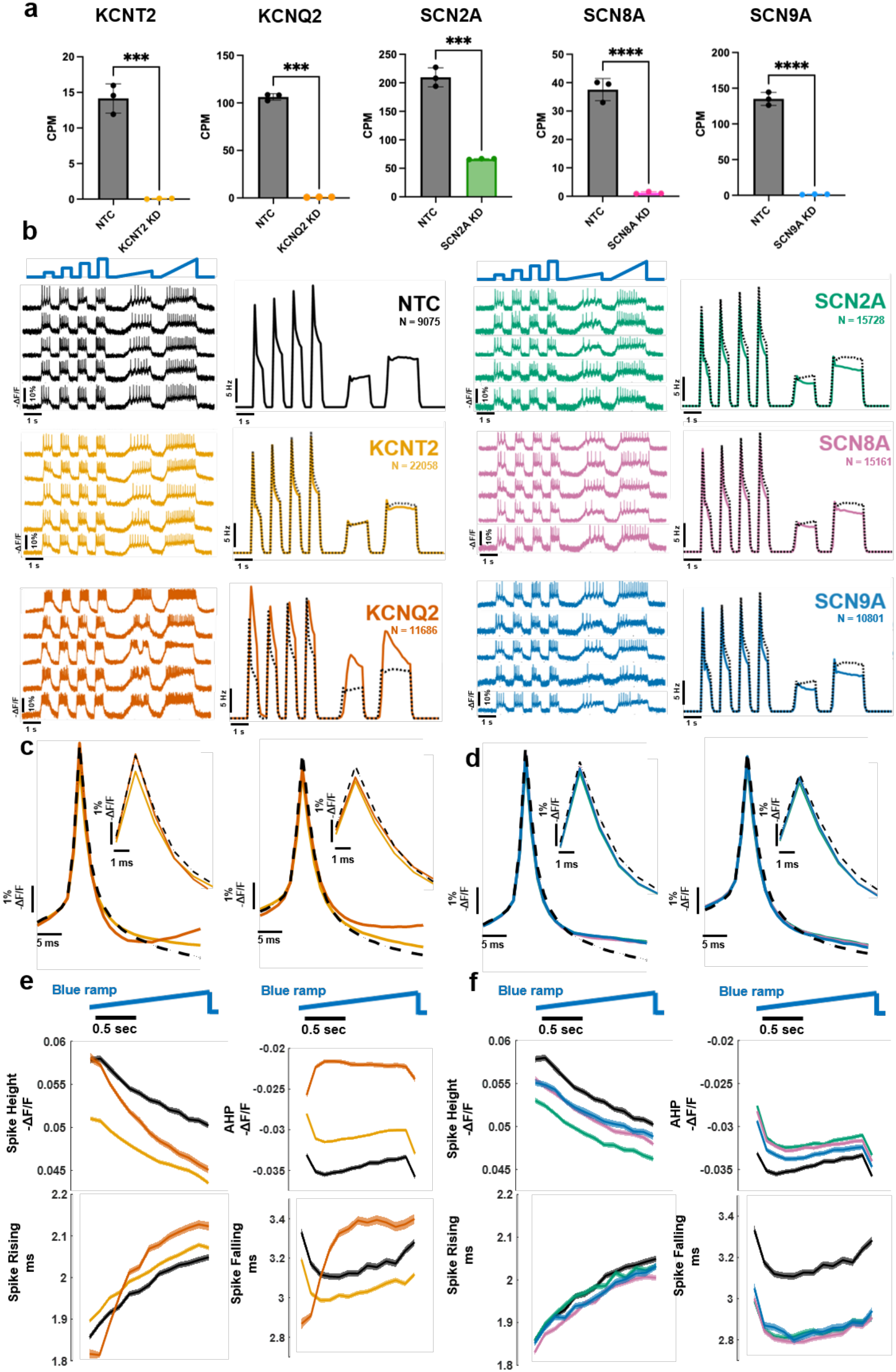
CRISPRi-mediated ion-channel knockdown produces gene-specific electrophysiological phenotypes. **(a)** Transcript abundance of each ion channel following CRISPRi-mediated knockdown with non-targeting control (NTC) cells shown for comparison. Transcript abundance is reported as counts per million (CPM) from bulk RNA seq. Statistical significance was assessed using an unpaired t-test. Error bars mean ± s.d. over independent rounds of differentiation. ***, 0.0001 < P ≤ 0.001; ****, P < 0.0001. **(b)** Optopatch excitability curves following targeted gene knockdown. Left: representative single-neuron voltage traces. Right: population-average firing-rates. NTC, N = 9,075; KCNT2, N = 22,058; KCNQ2, N = 11,686; SCN2A, N = 15,728; SCN8A, N = 15,161; and SCN9A, N = 10,801 cells. **(c, d)** Representative action potential waveforms for **(c)** potassium-channel and **(d)** sodium-channel knockdowns, measured during early and late portions of the ramp stimulation. The black dashed traces indicate NTC cells. Insets show an expanded view of the action-potential waveform. **(e, f)** Action-potential height, afterhyperpolarization (AHP), and spike rising and falling phase durations during the ramp stimulation for **(e)** potassium channel and **(f)** sodium channel knockdowns. NTC data are shown in both panels for reference. Shaded regions indicate SEM.

We recorded thousands of neurons for each perturbation using a standardized sequence of step and ramp blue-light stimuli (Fig. S8). The perturbations produced distinct effects on stimulus-evoked firing (Fig. 3b). KCNQ2 knockdown markedly increased firing across the stimulation protocol, consistent with the established role of the KCNQ2-mediated M-current in limiting neuronal excitability^32^. Knockdown of SCN2A, SCN8A, or SCN9A reduced evoked firing, most strongly late in sustained ramp and step stimuli^33–35^. KCNT2 knockdown also modestly reduced firing, rather than increasing it as loss of a potassium conductance might suggest. A recent study similarly found reduced excitability following KCNT2 perturbation in uniformly perturbed NGN2 neurons, but observed the opposite phenotype when perturbed cells were co-cultured with 50% wild-type neurons^19^. This context dependence suggests that the effect of KCNT2 loss involves non-cell-autonomous influences from neighboring cells, and not only its direct role in repolarization. The platform thus resolves gene-specific changes in excitability across large populations of perturbed neurons.

High-speed voltage imaging further enabled us to resolve gene-specific changes in action-potential waveform and its evolution during sustained stimulation. We compared population-averaged action-potential waveforms early and late in the ramp stimulus and quantified spike height, afterhyperpolarization (AHP), rising time, and falling time throughout the ramp (Fig. 3c–f; Fig. S9a; Fig. S10). NTC neurons showed progressive waveform adaptation during sustained firing, including a gradual decrease in spike height and an increase in spike rising time.

The two potassium-channel perturbations produced distinct action-potential waveform phenotypes (Fig. 3c,e). KCNT2 knockdown reduced spike height and AHP throughout the ramp, while producing a modest increase in rise time and a decrease in fall time. KCNQ2 knockdown showed a different and strongly time-dependent phenotype. At the beginning of the ramp, KCNQ2-knockdown neurons generated action potentials with relatively short rise and fall times. During sustained firing, however, both phases progressively lengthened, accompanied by a pronounced decrease in spike height. By the later portion of the ramp, KCNQ2-knockdown neurons exhibited substantially slower action-potential kinetics than at ramp onset. Thus, although KCNT2 and KCNQ2 both encode potassium channels, their knockdown produced clearly distinguishable effects on action-potential shape and adaptation.

Knockdown of the sodium-channel genes SCN2A, SCN8A, and SCN9A also produced characteristic waveform changes (Fig. 3d,f). All three perturbations reduced spike height and AHP relative to NTC neurons and were associated with shorter action-potential rise and fall times, with the reduction being more pronounced for the falling phase. Despite these shared effects, the magnitude of the phenotype differed among the three genes. SCN2A knockdown produced the strongest reduction in spike height and AHP, whereas SCN8A and SCN9A knockdown produced larger changes in spike rise and fall kinetics. Thus, perturbation of different voltage-gated sodium-channel genes generated overlapping but distinguishable action-potential waveform signatures.

Genetic perturbation also altered subthreshold voltage dynamics. Following abrupt termination of blue-light stimulation, the membrane potential of KCNQ2-knockdown neurons returned to baseline more slowly than that of NTC neurons (Fig. S9b,c). Thus, KCNQ2 perturbation affected not only action-potential firing and waveform adaptation but also membrane-potential dynamics outside of spikes. Together, these measurements show that the platform captures multidimensional electrophysiological phenotypes associated with individual genetic perturbations, spanning excitability, firing dynamics, action-potential waveform, adaptation, and subthreshold voltage responses.

### Optopatch stable hiPSC line supports optical electrophysiology across diverse cell types

Because Optopatch expression in the ACTB-SLEEK line is driven from the endogenous ACTB locus rather than a cell-type-specific promoter, the same engineered line can support electrophysiological measurements across multiple differentiated lineages. We therefore tested whether the line could be extended beyond NGN2-induced glutamatergic neurons by differentiating it into several cell types with distinct electrical properties (Fig. 4). Using NGN2-facilitated differentiation protocols, we generated glutamatergic neurons, cholinergic motor neurons, and astrocytes. NGN2-induced glutamatergic neurons expressed the vesicular glutamate transporter VGLUT1, whereas motor neurons expressed the cholinergic marker choline acetyltransferase (ChAT) together with the neuronal marker STMN2 (Fig. 4a,c)^36^. Astrocytes expressed the glutamate transporter EAAT1, consistent with acquisition of an astrocytic identity (Fig. 4e)^37^. Following TMP induction, all three neural-lineage cell types showed robust JF_608_-labeled Voltron2 fluorescence together with the constitutive fluorescent reporter, indicating that degron-controlled Optopatch expression was retained across these differentiation programs.

**Figure 4.**
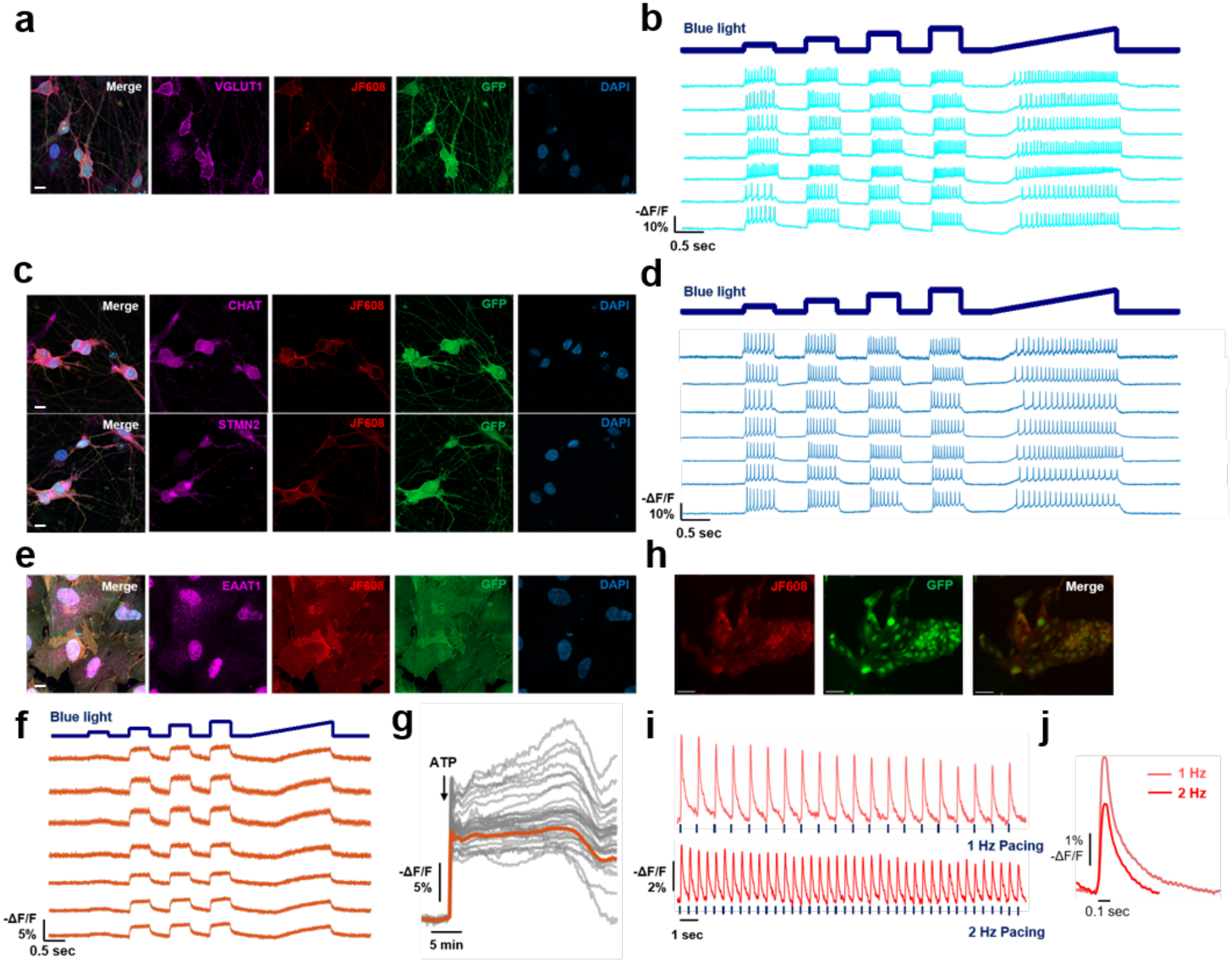
Optopatch hiPSC line enables optical electrophysiology across diverse human cell types. **(a)** Fluorescence images of NGN2-induced glutamatergic neurons. Magenta, glutamatergic marker VGLUT1; Red, JF_608_-labeled Voltron2; Green, GFP; Cyan, nuclei DAPI. Scale bar, 10 µm. **(b)** Voltage traces from NGN2-induced glutamatergic neurons. **(c)** Fluorescence images of NGN2-facilitated cholinergic motor neurons immunostained for choline acetyltransferase (ChAT; magenta, top) or stathmin-2 (STMN2; magenta, bottom). Red, JF_608_-labeled Voltron2; Green, GFP; Cyan, DAPI (nuclei). **(d)** Voltage traces from motor neurons. **(e)** Fluorescence images of NGN2-facilitated astrocytes immunostained for the astrocytic marker EAAT1 (magenta). Red, JF_608_-labeled Voltron2; Green, GFP; Cyan, DAPI (nuclei). **(f)** Voltage traces from astrocytes during blue-light stimulation. **(g)** Voltron2 recordings of astrocyte membrane-potential following 250 µM ATP application. Gray, individual cells; Orange, population mean. N = 180 cells. **(h)** Fluorescence images of iPSC-derived cardiomyocytes. Red, JF_608_-labeled Voltron2; Green, GFP. Scale Bar, 50 µm. **(i)** Voltage traces from cardiomyocytes during optical pacing at 1 Hz (top) and 2 Hz (bottom). Blue ticks indicate blue-light stimulation pulses. **(j)** Spike-triggered average waveforms during 1-Hz and 2-Hz optical pacing.

The resulting optical signals reflected the distinct electrophysiological properties of each cell type. Glutamatergic cortical neurons and cholinergic motor neurons generated robust trains of light-evoked action potentials during step and ramp blue-light stimulation (Fig. 4b,d). Astrocytes, in contrast, exhibited a qualitatively different electrical phenotype. Rather than generating fast action potentials, astrocytes showed slower membrane-potential responses to blue-light stimulation (Fig. 4f). We further tested whether Voltron2 could resolve chemically evoked astrocyte membrane dynamics by applying ATP, which activates purinergic signaling in astrocytes. ATP elicited prolonged and heterogeneous voltage responses across individual cells, including multiphasic changes that evolved over 30 minutes (Fig. 4g). These measurements demonstrate that the same reporter can capture both millisecond-scale action potentials in neurons and much slower membrane-potential dynamics in non-spiking cells.

To determine whether the ACTB-SLEEK strategy could also support a differentiation program independent of NGN2, we differentiated the same iPSC line into cardiomyocytes using a cardiac differentiation protocol^38^. Cardiomyocytes retained strong JF_608_-labeled Voltron2 and fluorescent-reporter expression after differentiation (Fig. 4h). Functionally, the cells generated clear cardiac action potentials and could be optically paced at both 1 and 2 Hz, with individual blue-light pulses reliably evoking action potentials at the corresponding stimulation frequency (Fig. 4i, **Movies 3, 4**). The resulting waveforms also exhibited frequency-dependent differences in action-potential shape (Fig. 4j), demonstrating that the platform can both modulate the activity of electrically excitable non-neuronal cells and resolve their electrophysiological responses.

Together, these results show that ACTB-targeted, TMP-gated Optopatch expression is not restricted to a single differentiation lineage or electrophysiological regime. The same engineered iPSC line supports measurements ranging from rapid action-potential firing in glutamatergic and motor neurons, to slower voltage responses in astrocytes, to optically paced cardiac action potentials. This cell-type portability substantially broadens the potential use of the platform for studying genetic and pharmacological effects on electrical phenotypes across diverse hiPSC-derived cell types.

## Discussion

We developed a platform for high-throughput optical electrophysiology and genetic perturbation in hiPSC-derived cells. The platform has three components: an engineered iPSC line combining inducible NGN2, dCas9-KRAB for CRISPRi, and TMP-gated Optopatch; an ultra-wide-field microscope that records optogenetically evoked activity from up to approximately 3,000 neurons in a 10-s acquisition; and an automated analysis pipeline that converts each movie into single-cell voltage traces and quantitative electrophysiological features. Engineering these components together addressed the expression, imaging, and analysis constraints that have limited the throughput of previous Optopatch experiments.

A central component is the stable ACTB-SLEEK Optopatch line. Targeting Optopatch to the endogenous ACTB locus under degron control resolved two competing requirements: healthy iPSCs during expansion, and strong, uniform Optopatch expression after differentiation. The ecDHFR(C12) degrons suppressed Voltron2 and CheRiff during iPSC maintenance, and trimethoprim restored expression before measurement. Expression from the ACTB locus remained stable throughout differentiation and was more uniform than expression from PiggyBac or lentiviral delivery. Reduced cell-to-cell variability is particularly valuable for genetic screening, where assay uniformity increases sensitivity to perturbation-induced effects and reduces the number of cells needed to resolve a phenotype.

The benchmark ion-channel perturbations demonstrate the value of combining large-scale measurements with information-rich voltage readouts. KCNT2 and KCNQ2 knockdown produced distinct waveform and adaptation signatures despite both genes encoding potassium channels, and the three sodium-channel knockdowns differed from one another in the magnitude of their effects on spike height and kinetics. Firing rate alone would not have separated these phenotypes. Simultaneous measurement of excitability, action-potential waveform, adaptation, and subthreshold voltage dynamics provides a multidimensional phenotype can distinguish perturbations that change overall activity through different underlying mechanisms.

The platform can be extended in several directions. Replacing dCas9-KRAB with a transcriptional activator such as dCas9-VPR would enable CRISPR activation (CRISPRa) and complementary gain-of-function screens^39^. The Optopatch cassette could be introduced into patient-derived or genome-edited disease lines, so that the electrophysiological consequences of disease-associated variants can be quantified across thousands of cells and screened against genetic modifiers or drugs. Because the ACTB locus drives expression in any lineage, we were able to record from iPSC-derived motor neurons, astrocytes, and cardiomyocytes as well as NGN2-induced excitatory neurons; the same strategy should apply to models of channelopathies, motor-neuron disease, and cardiac arrhythmia. TMP can also be added at the iPSC stage, allowing changes in membrane physiology to be followed through the transition from pluripotency to differentiated cell states.

Experimental design determines whether these measurements report intrinsic excitability or network activity. Here, we blocked synaptic transmission pharmacologically, so differences in firing reflected cell-autonomous electrophysiology. Synaptic transmission can instead be measured by separating stimulated presynaptic cells from voltage-imaged postsynaptic cells, either through patterned DMD stimulation or through genetic strategies that restrict the actuator and the indicator to mutually exclusive cell populations, as demonstrated with synOptopatch^40^.

The platform is also compatible with pooled genetic screening at substantially larger scale than demonstrated here. Because each cell’s phenotype is recorded together with its spatial position, pooled sgRNA libraries could be delivered to a single culture and perturbation identities recovered after imaging using optical pooled screening^42^. The CROP-seq-Multi-V2 architecture used here is compatible with such guide-recovery strategies^31^. Combining pooled perturbation, high-throughput voltage imaging, and spatial guide identification would map large genetic libraries onto multidimensional electrophysiological phenotypes. Similarly, the Optopatch measurements are compatible with diverse post-fixation measurements of gene expression, proteome, or morphology.

Optical electrophysiology nevertheless requires care in both experimental design and interpretation. Optogenetic stimulation introduces a light-dependent membrane conductance and is therefore not biophysically equivalent to direct current injection^43^, and prolonged or repeated activation of CheRiff can produce intracellular acidification through proton permeation of the channelrhodopsin^44^. Stimulation intensity and duration therefore merit attention. Fluorescence-based voltage imaging also reports relative rather than absolute membrane potential. The recently developed Voltage Measurement by Activated Photocycle (VMAP) approach exploits the photocycle dynamics of Voltron2 to provide an internally referenced measurement of absolute membrane voltage^41^, which could extend genetic screens to resting membrane potential and other slowly varying bioelectrical states.

Together, the engineered cell line, wide-field imaging system, and automated analysis pipeline provide a scalable framework for linking genetic perturbations to quantitative electrical phenotypes in human cells. Beyond throughput, the platform preserves information about action-potential shape, adaptation, subthreshold voltage dynamics, and stimulus-response relationships that are otherwise inaccessible in screening formats. These capabilities provide a path toward systematic functional-genomic maps of cellular electrophysiology across diverse human cell types and disease models.

## Supporting information

Movie 1: Wide-area Optopatch recording of NGN2 neurons

Movie 2: Wide-area Optopatch recording of NGN2 neurons

Movie 3: Wide-area Optopatch recording of hiPSC-derived cardiomyocytes, 1 Hz pacing

Movie 4: Wide-area Optopatch recording of hiPSC-derived cardiomyocytes, 2 Hz pacing

## Author contributions

J.Z. cloned the constructs, prepared the samples, acquired data and analyzed the data.

J.Z. and D.I. developed the video analysis pipeline.

J.Z., R.T.W. and P.C.B. developed tools for genetic perturbation in NGN2 neurons.

J.Z., A.M.B., T.U., M.W., R.N., E.F. and S.L.F. developed the stable Optopatch hiPSC line.

S.E.P. and L.D.L. provided HaloTag dyes and guidance on their use.

D.G. designed and built the ultrawide-field microscope.

D.I. developed Luminos bi-directional microscope control software.

K.S., H.F., and R.T.L. differentiated cardiomyocytes.

E.F. and R.N. supervised the work.

J.Z. and A.E.C. designed the project, oversaw execution, and wrote the manuscript.

## Acknowledgments

We thank Andrew Preecha, Camila Bodden, and Megan Hershfield for technical assistance. We thank Vy Ngo and Jax Xu at the Broad Institute, and Richard She in the laboratory of Jonathan Weissman, for assistance with establishing the Optopatch stable cell line and for sharing plasmids and related reagents. We thank Tsai-Wen Chen at National Yang Ming Chiao Tung University for sharing the Activity Localization Imaging (ALI) code prior to its publication. We thank the Harvard Center for Biological Imaging (RRID: SCR_018673) and the Harvard University Bauer Core Facility (RRID: SCR_022656) for infrastructure and technical support.

This work was supported by a Vannevar Bush Faculty Fellowship, the Harvard Brain Initiative, NIH grants R01NS126043, R01-NS133755, R01NS089076, R01MH128366, R01-HL174859, and R01-MH117042, and Chan Zuckerberg Initiative Dynamic Imaging Grant 2023-321177, the Simons Foundation Autism Research Initiative award (SFARI 890477), and The G. Harold and Leila Y. Mathers Charitable Foundation. T.U. was supported by the Huntington’s Disease Foundation. R.T.W. is supported by the National Science Foundation Graduate Research Fellowship Program under Grant No. 1745302.

## Data availability

Optopatch data recordings are available at: https://doi.org/10.7910/DVN/0Y3HEB

## Code availability

Data analysis code is available at: https://github.com/adamcohenlab/Zhang2026WideAreaOptopatchNGN2

Code contains functions for activity-based image segmentation, signal extraction, and assigning features to Optopatch traces.

## Optopatch hiPSC cell lines

The Optopatch hiPSC cell lines are available from AEC.

## Materials and Methods

### Genetic constructs

All constructs were generated using standard molecular cloning methods. Plasmid backbones were linearized by restriction enzyme digestion (New England Biolabs) and purified using the GeneJET Gel Extraction Kit (Thermo Fisher Scientific). DNA inserts were amplified using custom primers (Integrated DNA Technologies) and Q5 High-Fidelity DNA Polymerase (New England Biolabs) and assembled with linearized backbones using NEBuilder HiFi DNA Assembly Master Mix (New England Biolabs). All completed plasmids were verified by whole-plasmid sequencing (Plasmidsaurus).

#### Lentiviral Optopatch construct

The lentiviral Optopatch plasmid JZ026, encoding hSyn::Voltron2-ST-P2A-CheRiff-mEOS4a-ST, was generated by subcloning from the previously described JZ005: hSyn::Voltron2-ST-P2A-CheRiff-EGFP-ST construct (Addgene plasmid #258171). EGFP fused to CheRiff was replaced with the mEOS4a.

#### PiggyBac Optopatch construct

The PiggyBac donor vector TK4-EGFP and the corresponding transposase expression vector were gifts from Dr. Takeshi Uenaka and Dr. Marius Wernig, Stanford University. To generate the Optopatch PiggyBac donor plasmid JZ036: TK4-Optopatch-mNeon, the EF1α-driven puromycin-resistance cassette in the donor vector was replaced with an EF1α-driven mNeonGreen cassette to enable fluorescence-based selection. The CAG-driven EGFP expression cassette was replaced with the hSyn::Voltron2-ST-P2A-CheRiff-mEOS4a-ST cassette subcloned from JZ026.

#### CLYBL-targeted Optopatch construct

The TALEN nuclease plasmids targeting the CLYBL locus, pZT-C13-L1 and pZT-C13-R1, together with the CLYBL donor template pC13N_dCas9_BFP_KRAB (bsR), were gifts from Dr. Vy Ngo and Dr. Sami Farhi, Broad Institute. To generate the CLYBL-targeted Optopatch donor JZ037: pC13N_Optopatch (mNeon), the sequence between the left and right CLYBL TALEN homology arms was replaced with an expression cassette derived from JZ036. The donor contained an EF1α-driven mNeonGreen reporter and an hSyn::Voltron2-ST-P2A-CheRiff-mEOS4a-ST Optopatch cassette preceded by a ubiquitous chromatin-opening element (UCOE) and cHS4 insulator (full length). The entire expression cassette was also flanked by two cHS4 core insulator sequences to reduce transcriptional silencing.

#### ACTB-SLEEK degron-controlled Optopatch construct

The SLEEK donor template targeting the β-actin (ACTB) locus and the pX458 plasmid encoding SpCas9 together with an sgRNA targeting the ACTB locus were gifts from Richard She in the laboratory of Jonathan Weissman, Whitehead Institute. To generate a degron-regulated Optopatch cassette, ecDHFR(C12) destabilizing domains were first introduced into the JZ026 construct and fused to Voltron2 and CheRiff, respectively, generating a TMP-dependent Optopatch cassette. The cargo region of the ACTB-SLEEK donor was then replaced with this degron-regulated Optopatch cassette. An mNeonGreen fluorescent reporter was added downstream of the Optopatch components and linked by a P2A self-cleaving peptide, enabling FACS based picking of monoclonal line. The resulting donor construct JZ041: SLEEK-bACT-P2A-Optopatch-Degron-P2A-mNeon was used for SLEEK-mediated integration of degron-controlled Optopatch at the endogenous ACTB locus.

#### CROP-seq-Multi-V2 sgRNA constructs

The CROP-seq-Multi-V2 (CSMv2) template plasmid pRW1481 was a gift from Russell Walton and Dr. Paul Blainey (Broad Institute). For CRISPRi perturbations, sgRNA spacer sequences targeting each gene, together with non-targeting control sequences, were designed using CRISPick (Broad Institute). Spacer-specific insertion sequences were incorporated into dual-guide insert templates, which were synthesized by Twist Bioscience. Dual-guide inserts were cloned into the pRW1481 backbone by Golden Gate assembly (New England Biolabs) using the following thermal cycling program: 30 cycles of 42°C for 1 min and 16°C for 1 min, followed by 60°C for 5 min. Assembly products were transformed into NEB Stable chemically competent *E. coli* (New England Biolabs), and plasmid DNA was subsequently purified for downstream use.

### Cell culture

#### HEK233T cell culture

HEK293T cells were maintained in tissue-culture dishes (Corning) at 37°C and 5% CO₂ in Dulbecco’s Modified Eagle Medium (DMEM) supplemented with 10% fetal bovine serum (FBS), 1% GlutaMAX (Gibco, 35050061), 100 U/mL penicillin, and 100 µg/mL streptomycin (Gibco, 15140-122). Cells were passaged when cultures reached approximately 80% confluency. For passaging, cells were dissociated with 0.25% trypsin, and digestion was terminated by addition of four volumes of complete culture medium. Cells were collected by centrifugation at 300 × g for 5 min, resuspended in fresh culture medium, and replated at a 1:10 split ratio.

#### Lentivirus packaging

HEK293T cells were cultured in 35-mm tissue-culture dishes and transfected at approximately 80% confluency. For each dish, cells were transfected with 0.65 µg psPAX2 packaging plasmid, 0.30 µg VSV-G envelope plasmid, and 1.0 µg lentiviral transfer plasmid using TransIT-293 Transfection Reagent (Mirus Bio, MIR 2704). Virus-containing culture medium was collected at 48 and 72 h after transfection. The harvested medium was centrifuged at 500 × g for 5 min to remove cellular debris, and the clarified supernatant was concentrated approximately 10-fold using Lenti-X Concentrator (Takara Bio, 631231).

#### Mouse primary astrocyte culture

Cryopreserved primary P0 mouse astrocytes (ScienCell Research Laboratories, M1800-57) were maintained in Astrocyte Medium-animal (AM-a; ScienCell Research Laboratories, 1831). Culture flasks were coated overnight with poly-D-lysine (PDL) at 2 µg/cm² before cell seeding. For culture initiation, cryopreserved astrocytes were thawed rapidly in a 37°C water bath and diluted in prewarmed AM-a medium. Cells were then seeded into PDL-coated T75 flasks. When cultures reached approximately 90% confluency, astrocytes were dissociated using diluted trypsin consisting of 10% 0.25% trypsin-EDTA (Gibco, 25200056) and 90% 1× DPBS (Sigma-Aldrich, D8537). Trypsinization was terminated by addition of culture medium. Astrocytes were routinely expanded at a 1:3 to 1:5 split ratio. Cells were cryopreserved at passage 4 using CryoStor CS10 (STEMCELL Technologies, 07930).

#### CRISPRi lentiviral transduction and selection

On Day 1, ACTB-SLEEK iPSCs were seeded in 6-well plates at 31,250 cells/cm². On Day 2, cells were transduced in an arrayed format with concentrated CROP-seq-Multi v2 lentivirus at 1 µL virus per 200 µL mTeSR1 medium (STEMCELL Technologies, 85850). On Day 3, the virus-containing medium was replaced with standard iPSC culture medium. From Days 4–6, cells were selected with 5 µg/mL puromycin (Gibco, A1113803) for 72 h. After selection, cells were maintained in medium containing 0.5 µg/mL puromycin to allow recovery and for long-term maintenance of the transduced population.

#### Human iPSC culture

Culture plates were coated overnight with Geltrex (Gibco, A1413301) diluted 1:100 in DMEM/F12 (Gibco, 11320033). Human iPSCs were maintained on Geltrex-coated plates in mTeSR1 medium (STEMCELL Technologies, 85850) at 37°C and 5% CO₂, with medium replaced daily.

When cultures reached 80–90% confluency, cells were dissociated with Accutase (STEMCELL Technologies, 07920). Accutase was quenched by fivefold dilution with mTeSR1, and cells were collected by centrifugation at 300 × g for 5 min. The supernatant was removed, and cells were resuspended in mTeSR1 supplemented with 10 µM Y-27632 dihydrochloride ROCK inhibitor (STEMCELL Technologies, 72308). Cells were replated onto Geltrex-coated plates at a 1:20 split ratio. For the first 24 h after passaging, cells were maintained in mTeSR1 containing 10 µM Y-27632; thereafter, the medium was replaced with standard mTeSR1.

### Optopatch stable line generation

WTC11 iPSCs carrying a single copy of doxycycline-inducible mouse NGN2 at the AAVS1 locus and a single copy of CAG-driven dCas9-KRAB at the CLYBL locus were used as the parental line for subsequent genetic engineering.

To generate the CLYBL-targeted Optopatch line, iPSCs were co-transfected with the JZ037 donor plasmid, pC13N_Optopatch-mNeon, together with the CLYBL-targeting TALEN plasmids pZT-C13-L1 and pZT-C13-R1, using Lipofectamine Stem Transfection Reagent (Thermo Fisher Scientific, STEM00003). To generate the PiggyBac Optopatch line, iPSCs were co-transfected with the JZ036 donor plasmid, TK4-Optopatch-mNeon, and the PiggyBac transposase expression plasmid using Lipofectamine Stem. To generate the ACTB-SLEEK Optopatch line, iPSCs were co-transfected with the JZ041 donor plasmid, SLEEK-bACT-P2A-Optopatch-Degron-P2A-mNeon, and pX458 encoding SpCas9 together with an sgRNA targeting the ACTB locus, using Lipofectamine Stem (ThermoFisher, STEM00001).

Following transfection, cells were maintained for 10 days before fluorescence-based selection to minimize contribution from transient plasmid expression. At this time point, persistent mNeonGreen fluorescence was used as a marker for stable integration of the donor cassette. mNeonGreen-positive cells were isolated by fluorescence-activated cell sorting (FACS). For the PiggyBac line, sorted mNeonGreen-positive cells were collected and expanded as a polyclonal population. For the CLYBL-targeted and ACTB-SLEEK lines, sorted cells were plated at low density to enable isolation of individual colonies by direct visualization using an inverted microscope. Individual colonies were picked and expanded separately to establish monoclonal lines. Sorted cells were initially recovered in eTeSR medium (STEMCELL Technologies, 100-1215) and maintained until sufficiently expanded for passaging or colony isolation.

### NGN2 neuronal differentiation

For Optopatch imaging, glass-bottom imaging plates (Cellvis, P6-1.5H-N, P12-1.5H-N, and P96-1.5H-N) were coated overnight with poly-D-lysine (PDL; Gibco, A3890401) at 37°C, followed by an additional overnight coating with Geltrex (Gibco, A1413301) diluted 1:100 in DMEM/F12 (Gibco, 11320033) at 37°C. Primary mouse astrocytes were seeded onto the coated imaging plates at 24,000 cells/cm² and maintained for 3–6 days before addition of differentiating neuronal cells.

#### NGN2-induced excitatory neuron differentiation

For differentiation into excitatory neurons^28^, iPSCs were dissociated and seeded onto Geltrex-coated culture plates at 30,000 cells/cm². From Days 1–3, cells were maintained in induction medium consisting of DMEM/F12 (Gibco, 11320033), 1× N2 supplement (Gibco, 17502048), 1× GlutaMAX (Gibco, 35050079), and 0.3% glucose. On Day 1, the medium was supplemented with 2 µg/mL doxycycline (Sigma-Aldrich, D9891), 10 µM SB431542 (STEMCELL Technologies, 100-1051), 100 nM LDN193189 (STEMCELL Technologies, 72147), 2 µM XAV939 (STEMCELL Technologies, 100-1052), and 10 µM Y-27632 ROCK inhibitor (STEMCELL Technologies, 72308). On Days 2–3, cells were maintained in the same medium containing doxycycline, SB431542, LDN193189, and XAV939, but without ROCK inhibitor.

On Day 4, cells were dissociated using Accutase (STEMCELL Technologies, 07920) and replated onto astrocyte-containing imaging plates at 40,000 cells/cm². Beginning on Day 4, cells were maintained in neuronal medium consisting of BrainPhys Neuronal Medium supplemented with SM1 (STEMCELL Technologies, 05792), 1× GlutaMAX (Gibco, 35050079), 0.1 mM non-essential amino acids (Gibco, 11140076), and 0.3% glucose. The medium was further supplemented with 2 µg/mL doxycycline (Sigma-Aldrich, D9891), 10 ng/mL BDNF (RCD Systems, 248-BDB-250/CF), 10 ng/mL CNTF (RCD Systems, 257-NT-050/CF), and 10 ng/mL GDNF (RCD Systems, 212-GD-050/CF).

From Days 5–10, 1 µM floxuridine (Sigma-Aldrich, 1271008) and 1 µM uridine (Sigma-Aldrich, U3003) were added to suppress proliferating cells. Complete medium changes were performed on Days 5 and 10, with a half-medium change on Day 7. After Day 10, half of the medium was replaced every 2–3 days with fresh neuronal medium supplemented with doxycycline, BDNF, CNTF, and GDNF as described above.

#### NGN2-induced cholinergic motor neuron differentiation

For cholinergic motor neuron differentiation^36^, iPSCs were dissociated and seeded onto Geltrex-coated culture plates at 30,000 cells/cm². From Days 1–3, cells were maintained in DMEM/F12 (Gibco, 11320033) supplemented with 1× N2 (Gibco, 17502048), 1× GlutaMAX (Gibco, 35050079), and 0.3% glucose.

On Day 1, the induction medium was supplemented with 2 µg/mL doxycycline (Sigma-Aldrich, D9891), 10 µM SB431542 (STEMCELL Technologies, 100-1051), 100 nM LDN193189 (STEMCELL Technologies, 72147), 2 µM retinoic acid (Sigma-Aldrich, CAS 302-79-4), 2 µM Smoothened agonist (SAG; MedChemExpress, HY-12848C), and 10 µM Y-27632 ROCK inhibitor (STEMCELL Technologies, 72308). On Days 2–3, cells were maintained in the same induction medium containing doxycycline, SB431542, LDN193189, retinoic acid, and SAG, but without ROCK inhibitor.

From Days 4–6, cells were maintained in neuronal medium consisting of BrainPhys Neuronal Medium supplemented with SM1 (STEMCELL Technologies, 05792), 0.5× N2 supplement (Gibco, 17502048), 1× GlutaMAX (Gibco, 35050079), 0.1 mM non-essential amino acids (Gibco, 11140076), and 0.3% glucose. The medium was further supplemented with 2 µg/mL doxycycline, 2 µM retinoic acid, 2 µM SAG, 10 ng/mL BDNF, 10 ng/mL CNTF, and 10 ng/mL GDNF.

On Day 7, cells were dissociated using Accutase (STEMCELL Technologies, 07920) and replated onto astrocyte-containing imaging plates at 40,000 cells/cm². From Day 7 onward, cells were maintained in BrainPhys neuronal medium supplemented with SM1, 0.5× N2, 1× GlutaMAX, 0.1 mM non-essential amino acids, and 0.3% glucose, together with 2 µg/mL doxycycline, 10 ng/mL BDNF, 10 ng/mL CNTF, and 10 ng/mL GDNF.

From Days 7–10, 1 µM floxuridine and 1 µM uridine were added to suppress proliferating cells. After Day 10, half of the medium was replaced every 2–3 days with fresh neuronal medium supplemented with doxycycline, BDNF, CNTF, and GDNF as described above.

#### NGN2-induced astrocyte differentiation

For astrocyte differentiation^37^, iPSCs were dissociated and seeded onto Geltrex-coated culture plates at 30,000 cells/cm². On Day 1, cells were maintained in DMEM/F12 (Gibco, 11320033) supplemented with 2 µg/mL doxycycline (Sigma-Aldrich, D9891), 10 µM SB431542 (STEMCELL Technologies, 100-1051), 100 nM LDN193189 (STEMCELL Technologies, 72147), 2 µM XAV939 (STEMCELL Technologies, 100-1052), and 10 µM Y-27632 ROCK inhibitor. On Day 2, cells were maintained in DMEM/F12 supplemented with 2 µg/mL doxycycline, 5 µM SB431542, 50 nM LDN193189, and 1 µM XAV939, without ROCK inhibitor.

After Day 2, induced neural progenitor-like cells were dissociated with Accutase (STEMCELL Technologies, 07920) and replated onto Geltrex-coated plates at 15,000 cells/cm² in Astrocyte Medium (ScienCell, 1801) supplemented with 10 µM Y-27632 for the first 24 h after replating. Thereafter, cells were maintained in Astrocyte Medium without ROCK inhibitor for more than 30 days to promote astrocytic differentiation and maturation.

### Cardiomyocyte differentiation^38^

Three days before differentiation (Day −3), iPSCs were passaged and seeded into 6-well plates at 200k cells per well. Cells were maintained in StemFlex (Thermo A3349401), with complete medium changes on Days −2 and −1. On Day 0, differentiation was initiated when cultures reached approximately 70–85% confluency. Cells were switched to RPMI 1640 supplemented with B27 without insulin [RPMI/B27(−)] containing 7.5 µM CHIR99021. Cells were maintained under these conditions for 48 h at 37°C and 5% CO₂.

On Day 2, the medium was replaced with RPMI/B27(−) containing 5 µM Wnt inhibitor IWP4. The timing of this medium change was maintained as close as possible to 48 h after CHIR99021 addition. Cells were incubated for an additional 48 h. On Day 4, the medium was replaced with RPMI/B27(−) without small-molecule inhibitors. On Day 6, cells were switched to RPMI 1640 supplemented with B27 containing insulin [RPMI/B27(+)]. Thereafter, cultures were maintained in RPMI/B27(+) with medium changes every 48 h. On Day 11, cultures were subjected to metabolic selection by replacing the medium with glucose-free RPMI 1640 supplemented with B27 containing insulin and 1% penicillin-streptomycin. Cells were maintained in glucose-free medium for 48 h. On Day 13, cultures were returned to RPMI/B27(+) and subsequently maintained with medium changes every 48 h. For Optopatch imaging, cardiomyocytes were replated onto imaging plates 8 days before recording.

### Optopatch induction

To induce Optopatch expression in the ACTB-SLEEK line, 10 µM trimethoprim (TMP; Cayman Chemical, CAS 738-70-5) was added beginning 7 days before imaging for standard mature-differentiation recordings. For examining electrophysiological maturation between Days 4 and 28, TMP was added 4–5 days before each imaging time point to allow sufficient Optopatch expression. For lentiviral Optopatch expression, neurons were transduced on Day 14 with concentrated lentivirus at 1 µL virus per 100 µL culture medium.

### Bulk RNA sequencing

ACTB-SLEEK line was transduced with CROP-seq-Multi-V2 lentiviral vectors expressing either non-targeting control sgRNAs or sgRNAs targeting specific ion channels, differentiated into NPCs and replated on coated glass bottom six-well plates at a density of 70,000 cells/cm² on differentiation day 4, without astrocyte co-culture. Cells were subsequently maintained under the neuronal differentiation and culture conditions described above. At Day 28, cells were dissociated using Accutase (STEMCELL Technologies, 07920) for 5 min. The dissociation reaction was quenched by adding five volumes of BrainPhys Neuronal Medium supplemented with SM1 (STEMCELL Technologies, 05792). Cell suspensions were centrifuged at 500 × g for 5 min, and the supernatant was carefully aspirated. Cell pellets were resuspended in DNA/RNA Shield (Zymo Research, R1100-250) for cell lysis and RNA preservation. Three independent wells were collected for each experimental condition (n = 3). Samples were submitted to Plasmidsaurus for bulk RNA sequencing.

### Immunocytochemistry

Cells were washed three times with PBS for 5 min, and fixed directly in culture plates by replacing the culture medium with 4% paraformaldehyde (PFA) in PBS and incubating for 20 min at room temperature. Fixed cells were washed three times with PBS for 5 min each and permeabilized with 0.5% Triton X-100 in PBS for 10 min at room temperature. Cells were then rinsed three times for 1 min each with wash buffer consisting of PBS containing 0.1% Triton X-100. Cells were blocked for 10 min at room temperature in antibody dilution buffer (AbDil) consisting of 2% bovine serum albumin (BSA) in wash buffer. Primary antibodies were diluted in AbDil and added to the cells at 200 µL per well, followed by incubation for 1 h at room temperature on a nutating mixer. Cells were subsequently washed three times for 5 min each with wash buffer. Fluorophore-conjugated secondary antibodies were diluted in AbDil and incubated with the cells for 1 h at room temperature on a nutating mixer. Cells were then washed three times for 5 min each with wash buffer. Nuclear counterstaining was performed using DAPI at 1 µg/mL in AbDil for 10 min at room temperature. Cells were washed three additional times with wash buffer for 5 min each, followed by two 5-min washes with PBS before imaging.

For lineage characterization, the following primary antibodies were used: rabbit polyclonal anti-EAAT1/SLC1A3 antibody (Boster Biological Technology, PA2185) for astrocytes; rabbit polyclonal anti-Stathmin-2/STMN2 antibody (Novus Biologicals, NBP1-49461) and goat polyclonal anti-choline acetyltransferase (ChAT) antibody (MilliporeSigma/Chemicon, AB144P) for cholinergic motor neurons. Rabbit primary antibodies were detected using donkey anti-rabbit IgG conjugated to Alexa Fluor 555 (Thermo Fisher Scientific/Fisher, A-31572), and the goat primary antibody was detected using anti-goat IgG (H+L) conjugated to CF750 (Sigma-Aldrich, SAB4600444).

### Imaging conditions

#### Imaging buffers

For NGN2-induced neurons and astrocytes, recordings were performed in XC imaging buffer containing 125 mM NaCl, 15 mM HEPES, 25 mM glucose, 2.5 mM KCl, 1 mM MgCl₂, and 2 mM CaCl₂. The buffer was adjusted to pH 7.3 before use. Synaptic blockers, 10 μM NBQX (Tocris 1262/10), 25 μM D(-)-2-amino-5-phosphonovaleric acid (Tocris 0106/10), and 20 μM gabazine (Tocris SR-95531) were added. For cardiomyocyte imaging, recordings were performed in a Tyrode-like cardiac imaging buffer containing 135 mM NaCl, 5.4 mM KCl, 1 mM MgCl₂·6H₂O, 1.5 mM CaCl₂·2H₂O, 0.33 mM NaH₂PO₄·H₂O, 5 mM D-glucose, and 5 mM HEPES. The buffer was adjusted to pH 7.3 before use.

#### JF_C08_ labeling and imaging preparation

For Voltron2 imaging, cells were incubated with 200 nM JF_608_ in culture medium for 30 min. Cultures were then washed twice and incubated in imaging buffer for 15 min. Immediately before recording, the buffer was replaced with fresh imaging buffer to minimize extracellular dye background.

#### Customized ultrawide-field microscope

Optopatch movies were acquired using a custom-built ultrawide-field microscope (Fig. 2a). Illumination was provided by a red LED (OSRAM, LE A P2MQ, 613 nm) for excitation of JF_608_-labeled Voltron2 and a blue LED (Luminus, PT-121-B, 460 nm) for activation of CheRiff. Red and blue illumination were spectrally filtered using 600/37-nm (IDEX, FF01-600/37-32) and 457/50-nm (IDEX, FF01-457/50-32) bandpass filters, respectively. Light from each LED was collected using a two-lens assembly (Thorlabs, LA4148-A and ACL3026U-A) and combined using a dichroic mirror (IDEX, FF580-FDi01). The combined illumination was passed through a homogenizer incorporated into a custom digital micromirror device (DMD) assembly (Digital Light Innovations, 3DLP9000), enabling spatial and temporal control of illumination patterns. Light reflected from the DMD was collected using a 105-mm camera lens (Sigma, ART 105 mm F1.4 DG HSM) and relayed to the sample through a 2× objective with 0.5 numerical aperture (Olympus, MVPLAPO 2XC). Unless otherwise specified, the LEDs were driven at 5 A, producing an illumination intensity of approximately 20 mW/mm² at the sample plane. Fluorescence emission was collected through the same objective and separated from excitation light using a custom dichroic mirror (Chroma, zt488/607tpc, 50 × 72 × 3 mm) together with an emission filter (IDEX, AF01-653/47). Fluorescence was projected onto a scientific CMOS camera (Teledyne Kinetix) using a second 105-mm camera lens (Sigma, ART 105 mm F1.4 DG HSM), yielding an effective magnification of 2.3×. Cultures were maintained using a stage-top incubator (Tokai Hit, INUBG2ATFP-WSKM) with temperature and humidity control.

#### Microscope control and acquisition synchronization

Microscope hardware control and data acquisition were performed using Luminos software^29^. Camera acquisition was synchronized using frame-trigger pulses generated by a National Instruments DAQ system (USB-6353) referenced to the DAQ internal 100-kHz clock. The same clock was used to synchronize camera acquisition with DMD pattern switching and LED drive waveforms, enabling precise temporal coordination of optogenetic stimulation and voltage imaging.

### Data analysis

#### Automated activity localization imaging and trace extraction

A custom MATLAB implementation of activity localization imaging (ALI)^30^ was used to identify active cells and to extract their fluorescence traces. Raw movies were first corrected for spatially varying background fluorescence. The field of view was then divided into small regions containing one or more cells using an intensity-based threshold. The fluorescence time-series at each pixel was subjected to a temporal high-pass median filter to emphasize spikes relative to slower fluctuations. Because Voltron2 fluorescence decreases upon membrane depolarization, the median filtered fluorescence changes were then inverted to make spikes appear positive-going. The fluorescence time course within each spatial region was averaged to generate a preliminary temporal trace. Candidate action potentials were then identified from these traces via a locally determined threshold.

For each detected action potential, we determined the centroid of the fluorescence change. Spike centroids were clustered according to spatial proximity – closely spaced centroids were assumed to originate from the same cell. For each cluster of spike events, we calculated a spike-triggered mean image of -ΔF, to generate an activity-defined spatial footprint of the cell. Cell-specific temporal signals were then recovered from the background-corrected raw fluorescence movie using pseudoinverse-based spatial unmixing. This procedure resolved signals from touching or partially overlapping neurons.

Extracted fluorescence traces were subsequently corrected for slow baseline drift and photobleaching by fitting the baseline fluorescence with a two-component exponential function. The measured fluorescence was normalized to the fitted baseline, and the polarity was inverted to calculate -ΔF/F, such that membrane depolarization corresponded to a positive-going voltage signal. The resulting normalized traces were used for subsequent spike detection and electrophysiological feature extraction.

#### Extraction of action-potential and subthreshold electrophysiological features

Action-potential waveform features were extracted from each action potential in each cell. Spike onset was defined as the frame corresponding to the maximum second derivative of the waveform during the rising phase. Spike height was defined as the difference between the action-potential peak and the voltage at spike onset (Fig. S9a). Spike width was measured between the rising and falling crossings at the level corresponding to 80% repolarization from the spike peak, with sub-frame interpolation. Spike rising time was defined as the interval between spike onset and the action-potential peak. Spike falling time was defined as the interval between the action-potential peak and the post-peak crossing of the 80%-repolarization level. Afterhyperpolarization (AHP) was quantified as the difference between the spike-onset voltage and the minimum voltage reached within a 37.5-ms window following the action-potential peak, with the search window truncated before the peak of the next detected spike, if present.

We analyzed the voltage trajectory immediately after blue-light offset to estimate a repolarization time-constant (Fig. S9b,c). Traces containing action potentials within the analyzed repolarization interval were excluded. The post-stimulus response was scaled to the range [1, 0] and fit with a single-exponential decay:

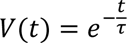

where τ is the exponential decay time constant.

