## Supplementary Information for "A high-throughput all-optical platform for mapping gene perturbations to electrophysiology"

Figures S1 – S10
Tables S1 – S2
Supplementary Movie captions

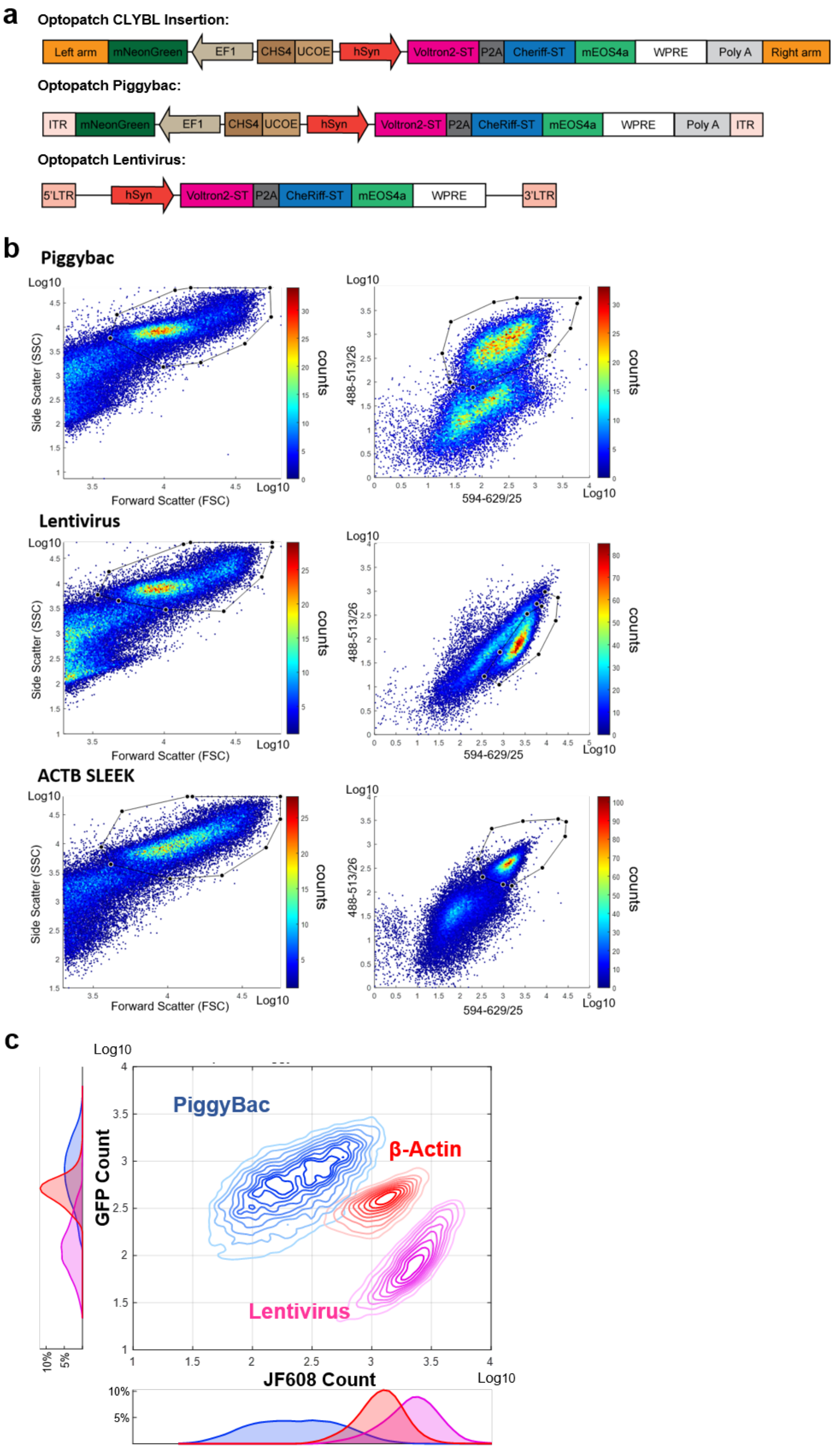


**Figure S1. Comparison of Optopatch expression strategies in iPSC-derived neurons.** (a) Schematics of Optopatch expression in DIV28 neurons using targeted CLYBL insertion, PiggyBac integration, and lentiviral delivery. (b) Representative flow-cytometry density plots showing forward- and side-scatter gating and fluorescence distributions for PiggyBac, lentiviral, and β-actin stable-line expression in DIV28 neurons. (c) Overlaid density contour plots comparing JF608 and GFP fluorescence across the three expression strategies. Contour lines mark 10-percentile increments in cell density.

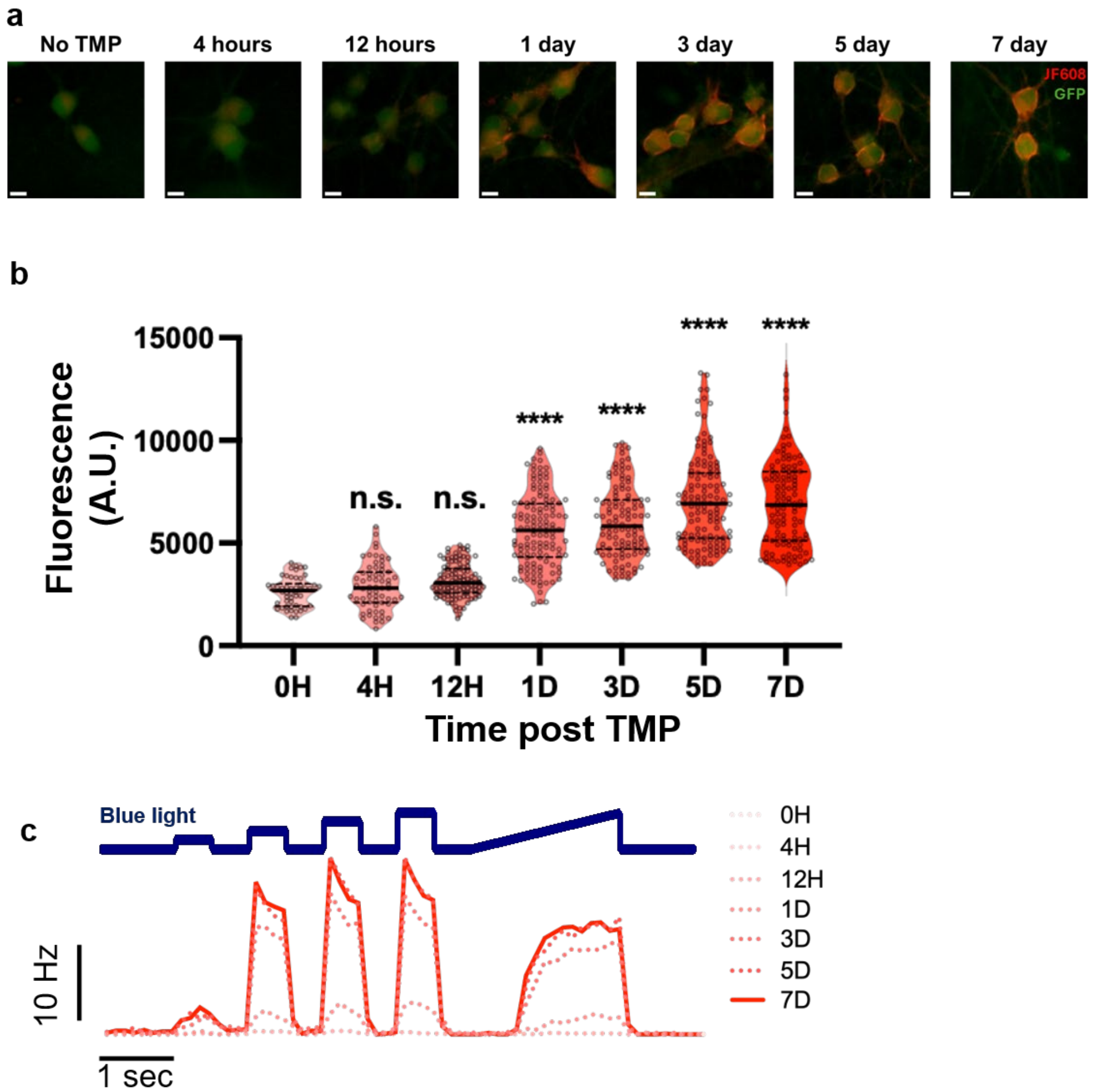


**Figure S2. Time-dependent induction of Optopatch expression and function following TMP treatment.** (a) Representative fluorescence images of neurons from the β-actin SLEEK stable Optopatch line without TMP treatment and with 4 h, 12 h, 1 day, 3 days, 5 days, and 7 days of TMP treatment. Neurons were imaged at DIV28. Scale bars, 10 µm. (b) Quantification of cellular fluorescence intensity over the TMP treatment time course. 0 h, n = 50; 4 h, n = 63; 12 h, n = 108; 1 day, n = 128; 3 days, n = 115; 5 days, n = 132; 7 days, n = 110 cells. Statistical significance was assessed by one-way ANOVA. n.s., $P > 0.05$; ****, $P < 0.0001$. (c) Representative population firing-rate responses to the blue-light stimulation protocol following different durations of TMP treatment. Same numbers of cells as in (b).

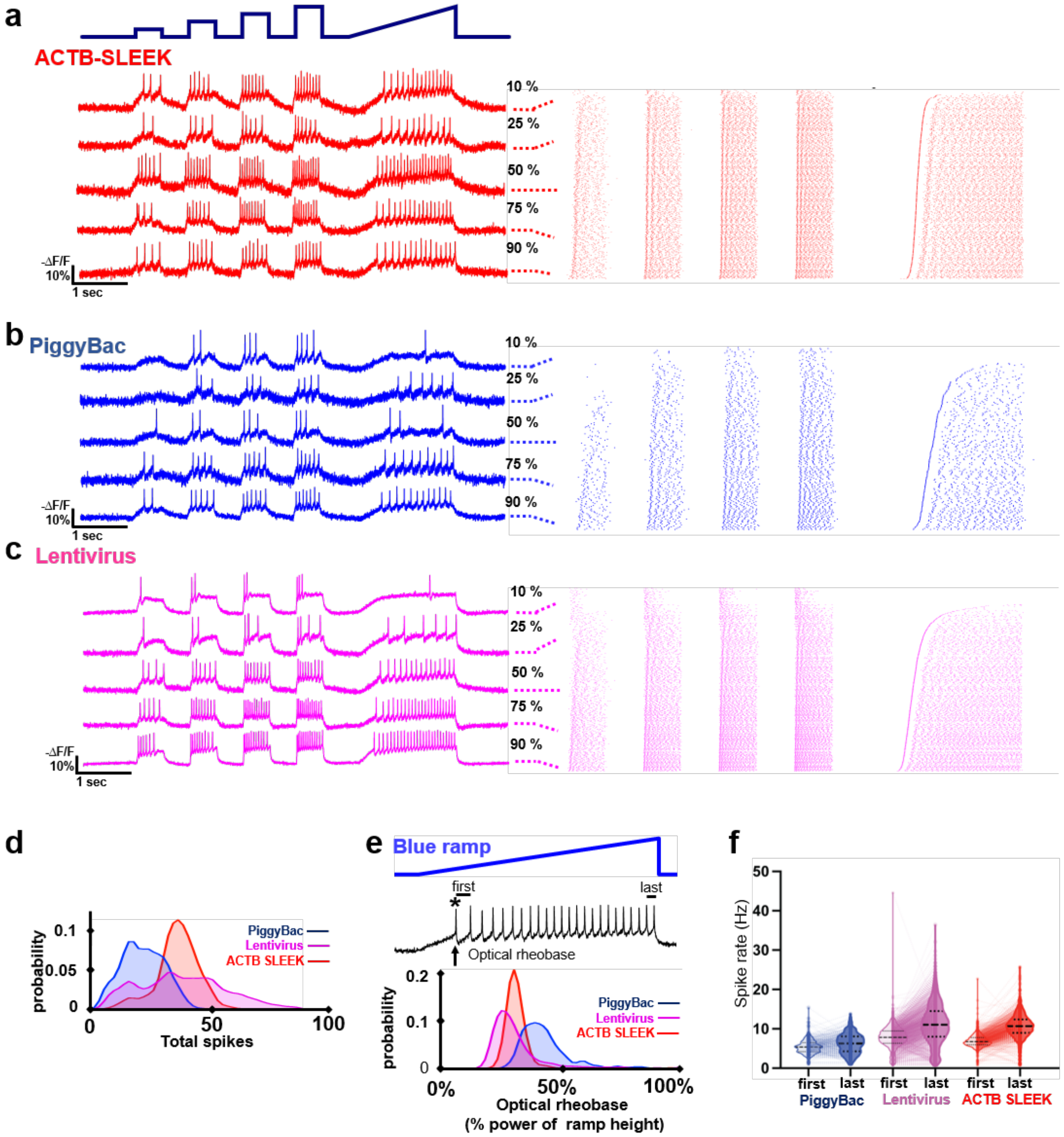


**Figure S3. Comparison of Optopatch performance in neurons expressing Optopatch by PiggyBac integration, lentiviral transduction, or ACTB-SLEEK integration.** (a-c) Left: representative voltage traces from neurons at the indicated percentiles of excitability. Right: raster plots of all imaged neurons during Optopatch step and ramp protocol, sorted by optical rheobase. ACTB-SLEEK (data repeated from Fig. 1g), N = 1,046 neurons; PiggyBac, N = 362 neurons; Lentivirus, N = 2,053 neurons. (d) Distribution of total optically evoked spikes per neuron during step and ramp protocol. (e) Top: Definition of electrophysiological parameters extracted from the blue-light ramp. Optical rheobase was defined as the blue-light intensity at the first evoked spike. Initial and final firing rates were calculated from the first and last interspike intervals, respectively. Bottom: Distribution of optical rheobase for PiggyBac, lentiviral, and ACTB-SLEEK neurons. (f) Initial and final firing rates during the blue-light ramp for the three Optopatch expression strategies. Thin lines connect measurements from individual neurons; violin plots show the population distributions. Thick and thin dashed lines indicate the median and interquartile range, respectively.

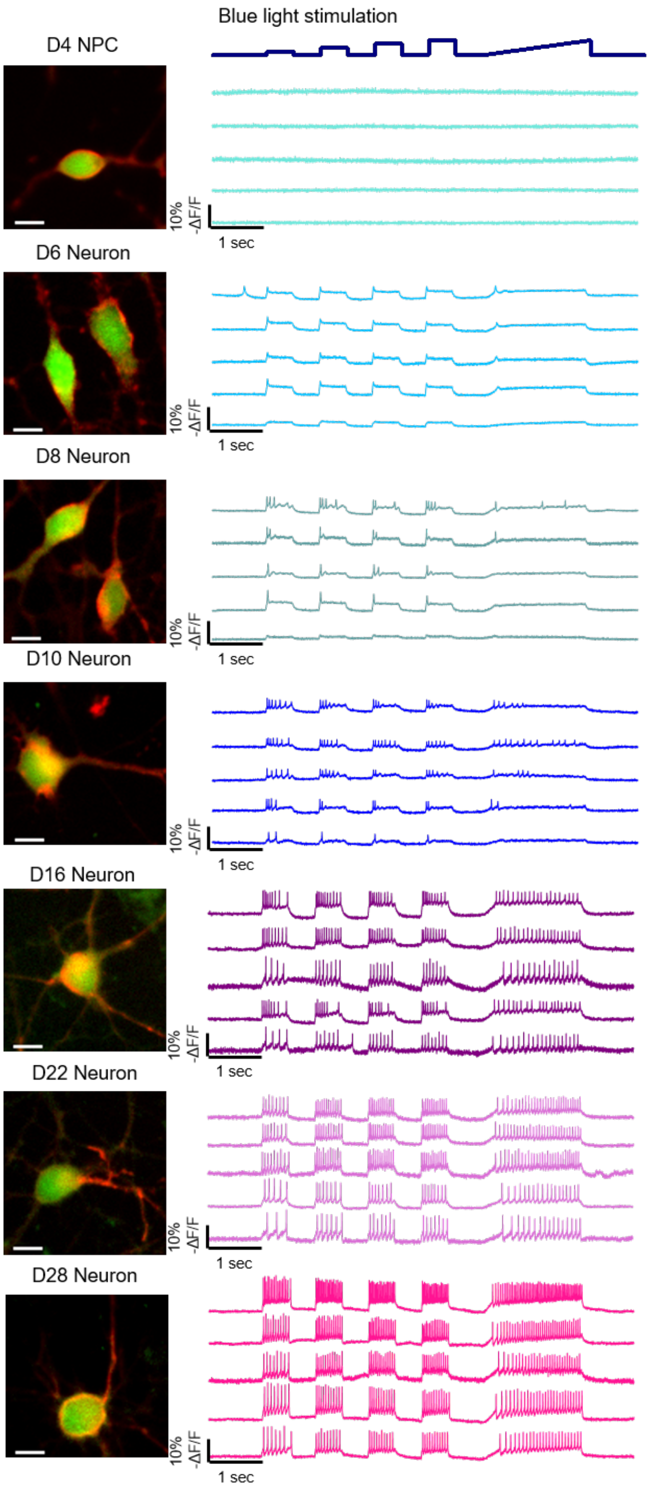


**Figure S4. Development of optically evoked electrophysiological responses during neuronal differentiation.** Representative fluorescence images and optical voltage traces from the β-actin SLEEK Optopatch line at the neural progenitor stage (D4 NPC) and at D6, D8, D10, D16, D22, and D28 of neuronal differentiation. Five representative cellular traces are shown for each stage. Scale bars, 10 μm.

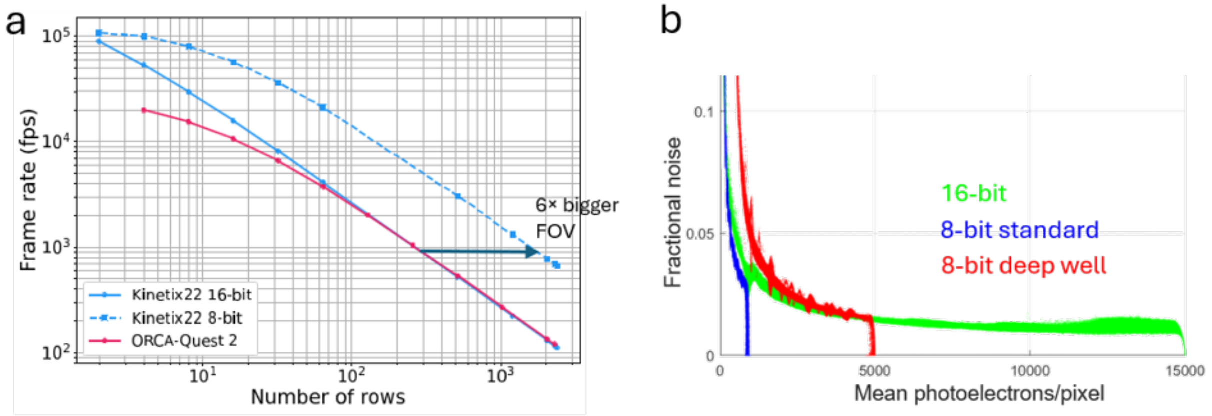


**Figure S5. Adjusting camera parameters to optimize FOV and SNR.** (a) 8-bit mode leads to 6-fold larger FOV at 1 kHz. (b) Deep-well mode (red) improves shot-noise-limited SNR at 8-bit readout relative to 8-bit standard mode (blue). For near-saturating signals, 8-bit deep-well mode and 16-bit mode have similar fractional noise.

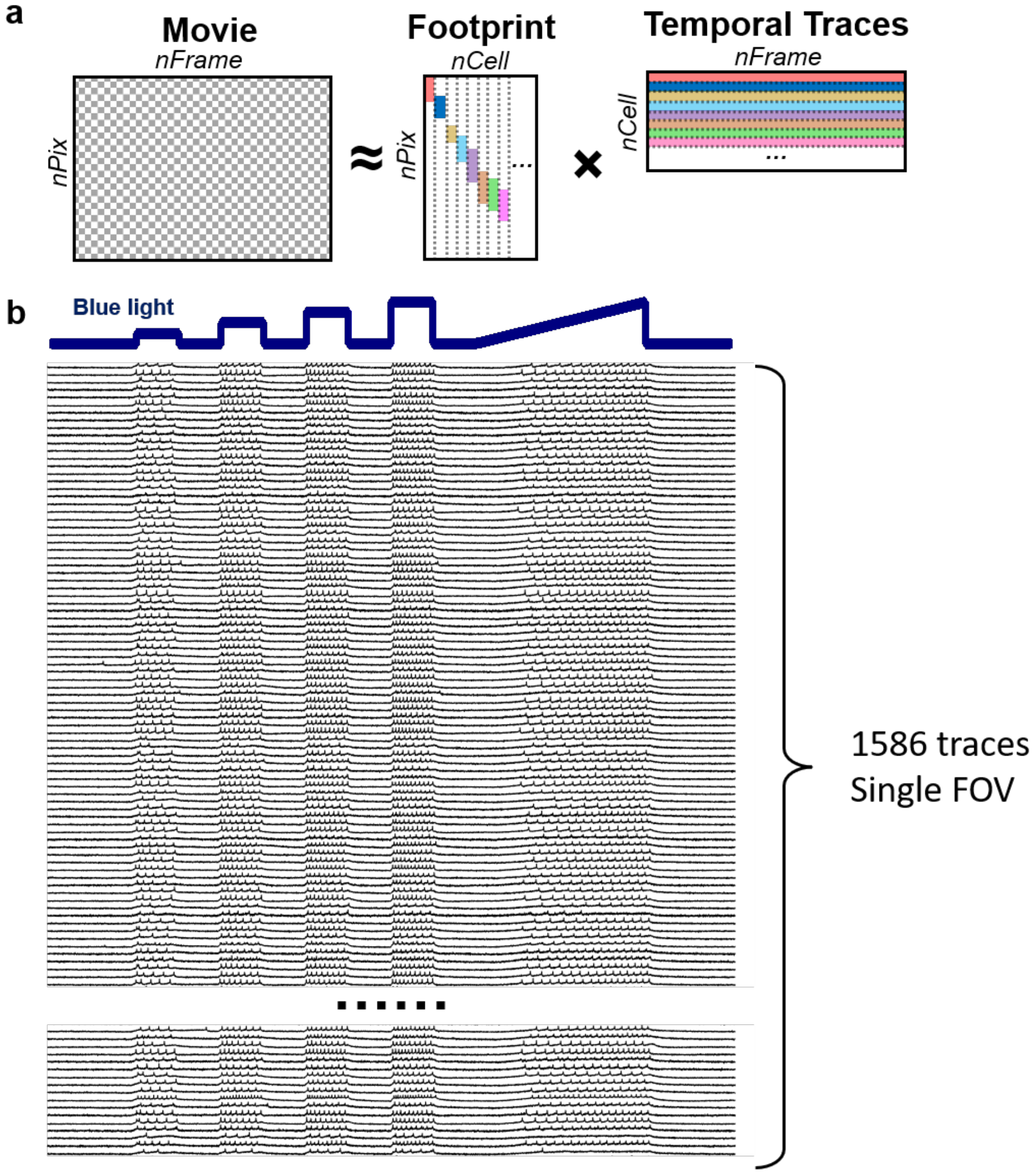


**Figure S6. Extraction of single-cell optical voltage traces from movies.** (a) Schematic of the matrix-factorization framework used to represent the fluorescence movie as the product of spatial cellular footprints and their corresponding temporal fluorescence traces. nPix is the number of pixels per frame; nFrame is the number of frames. (b) Extracted optical voltage traces from a randomly selected subset of 1,586 cells recorded within a single field of view.

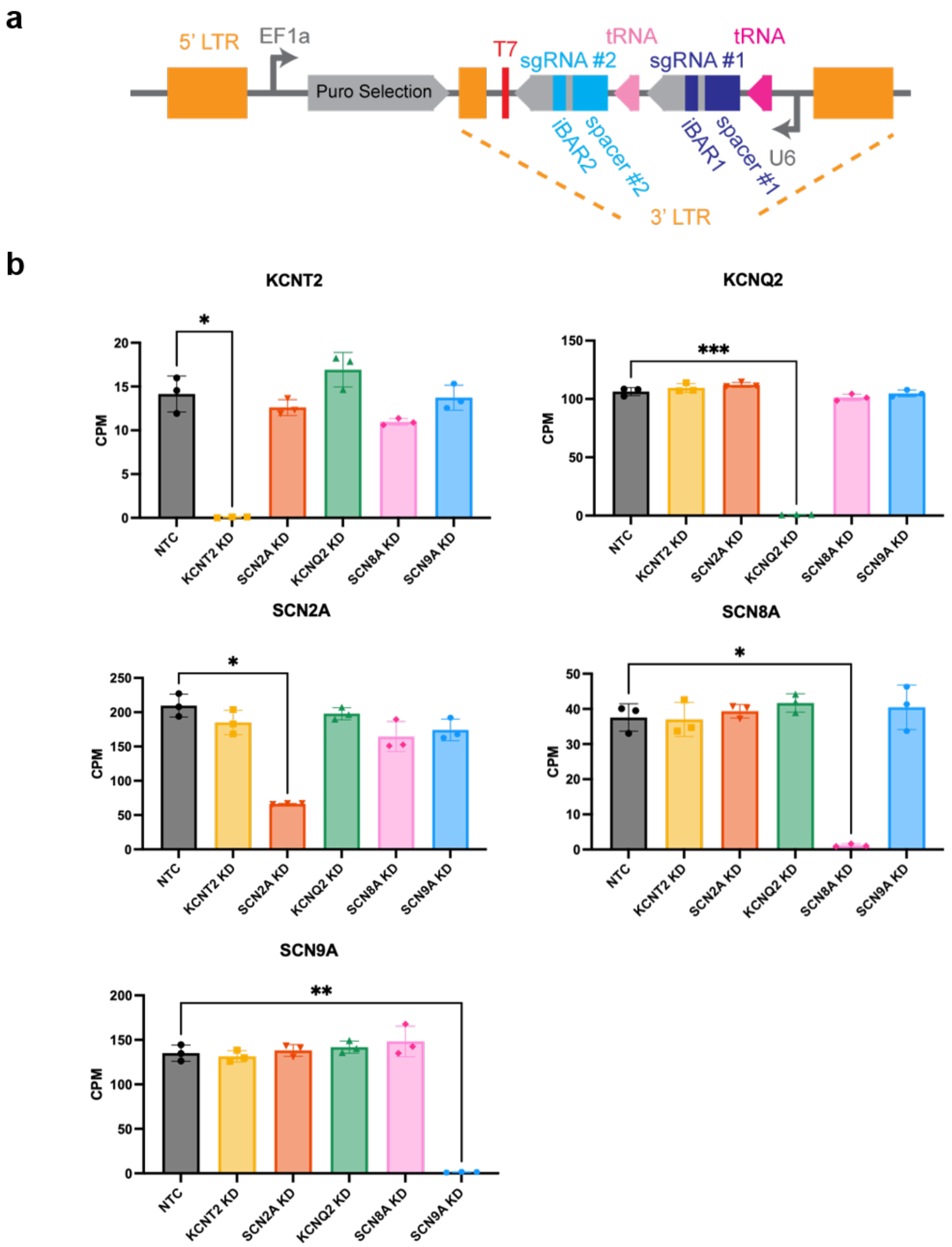


**Figure S7. CROPseq-multi-v2 vector design and validation of target-gene knockdown.** (a) Schematic of the CROPseq-multi-v2 lentiviral construct containing an EF1α-driven puromycin-selection cassette and two sgRNAs separated by tRNA elements. (b) Bulk RNA seq validation of CRISPRi-mediated knockdown of KCNT2, KCNQ2, SCN2A, SCN8A, and SCN9A. Transcript counts per million (CPM) are compared among non-targeting control (NTC), and knockdown. Symbols denote replicates from different dishes. Plots show mean ± s.d.. Statistical significance was assessed by Brown-Forsythe and Welch ANOVA tests. N.s. ($P > 0.05$) is not marked. *, $0.01 < P \leq 0.05$, **, $0.001 < P \leq 0.01$, ***, $0.0001 < P \leq 0.001$.

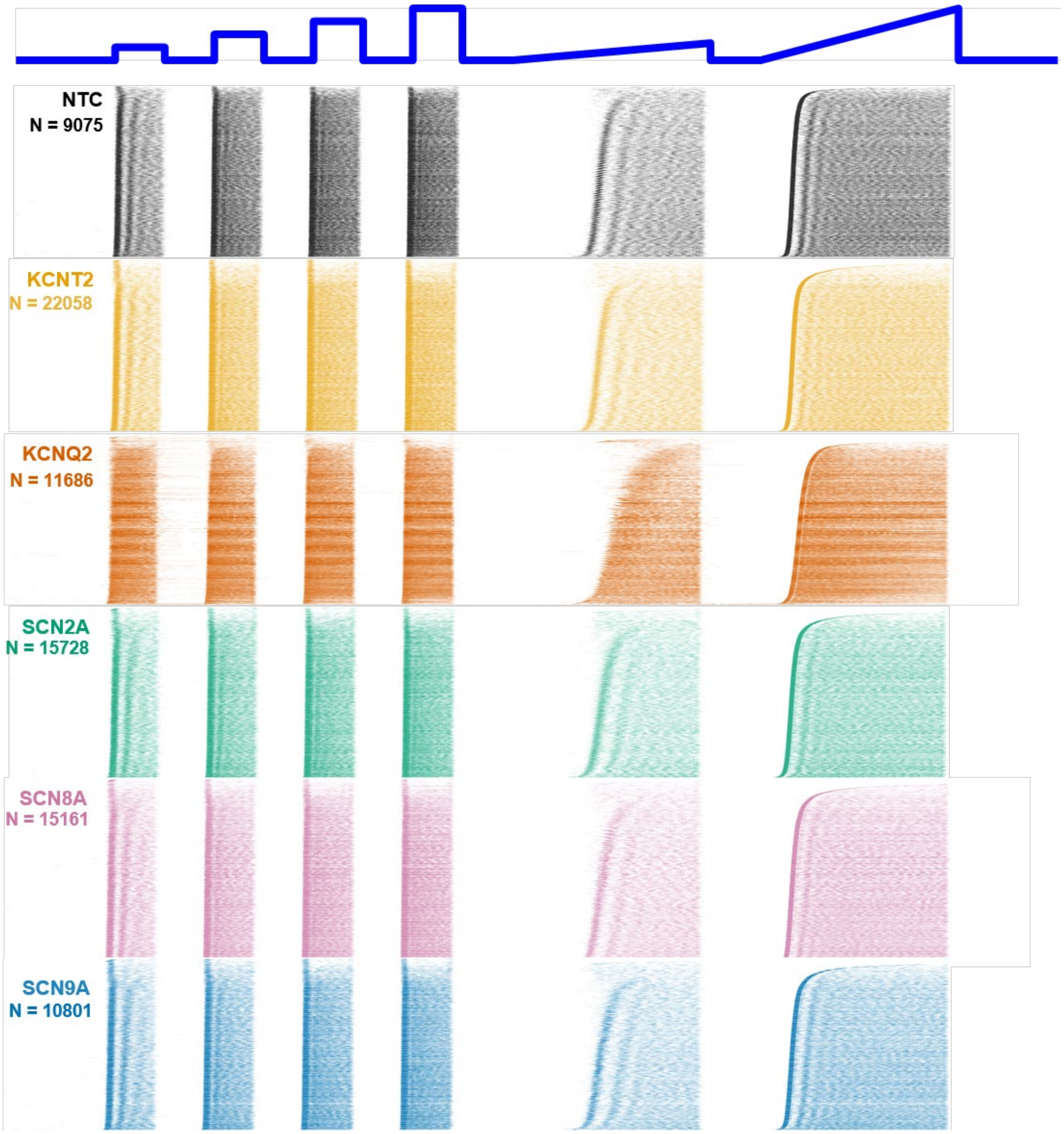


**Figure S8. Population-scale optical electrophysiology responses following ion-channel knockdown.** Raster plots of spiking activity from all imaged neurons transduced with non-targeting control sgRNAs (NTC; N = 9,075) or sgRNAs targeting KCNT2 (N = 22,058), KCNQ2 (N = 11,686), SCN2A (N = 15,728), SCN8A (N = 15,161), or SCN9A (N = 10,801). Each row represents the spiking events of an individual neuron in response to the blue-light stimulation protocol shown at the top. Neurons are sorted by optical rheobase measured during the second ramp stimulus.

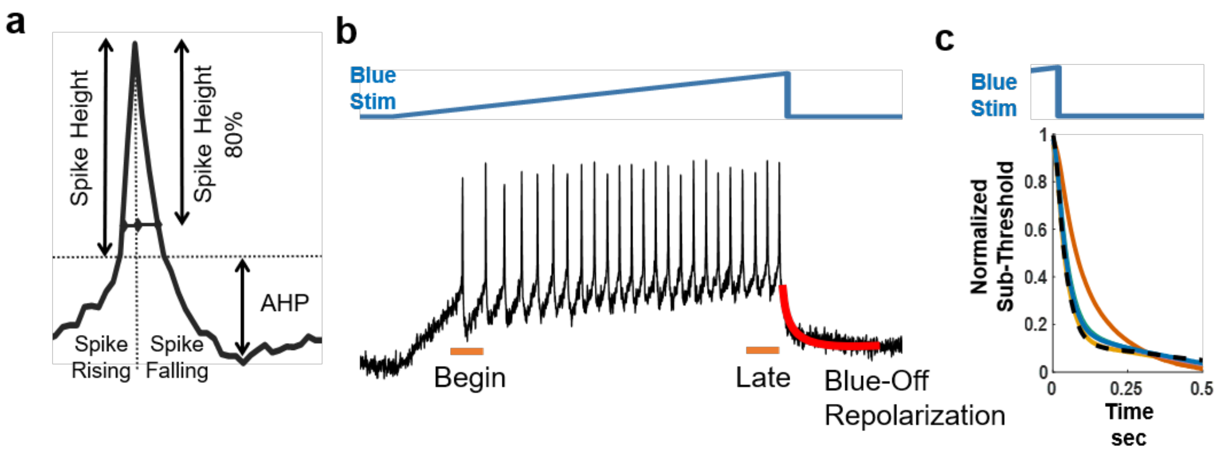


**Figure S9. Extraction of action-potential and subthreshold electrophysiological features from optical voltage traces.** (a) Schematic illustrating waveform features extracted from individual action potentials, including spike height, spike width, afterhyperpolarization (AHP), and rising and falling phases of the action potential. (b) Representative optical voltage response during a blue-light ramp stimulus. The early and late action potentials within the ramp are indicated, and the red segment highlights membrane repolarization following termination of blue-light stimulation. (c) Representative normalized subthreshold repolarization trajectories following blue-light offset.

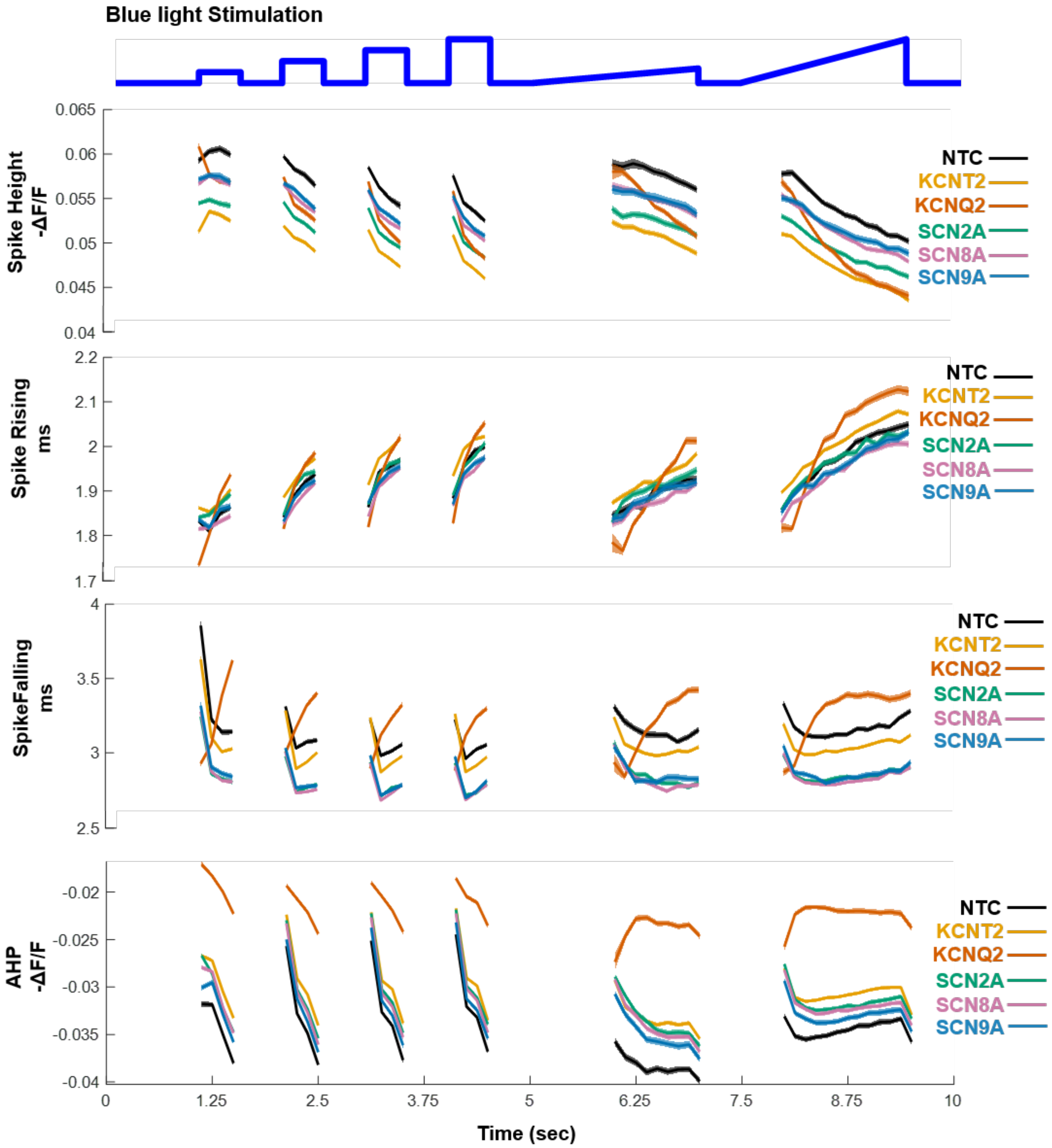


**Figure S10. Ion-channel perturbations alter action-potential waveform properties with distinct signatures.** Population trajectories of action-potential features during the step and ramp blue-light stimulation protocol for NTC neurons and neurons with CRISPRi knockdown of KCNT2, KCNQ2, SCN2A, SCN8A, or SCN9A. From top to bottom, plots show spike height (−ΔF/F), spike rising phase (ms), falling phase (ms), and AHP (−ΔF/F). Measurements are plotted across successive stimulation epochs to reveal perturbation-dependent differences in action-potential waveform and their evolution during sustained firing. Cell numbers are in Fig. S8. Error bar, SEM.

| | Group | Mean | Median | Std | Q1 | Q3 | N |
|---|---|---|---|---|---|---|---|
| JF608 (A.U.) | PiggyBac | 326.02 | 229.51 | 293.57 | 119.71 | 440.09 | 15956 |
| | Lentivirus | 2419.89 | 2205.31 | 1280.02 | 1532.74 | 3044.31 | 12548 |
| | ACTB SLEEK | 1304.89 | 1204.00 | 677.04 | 888.04 | 1581.62 | 10962 |
| GFP (A.U.) | PiggyBac | 752.93 | 582.48 | 595.46 | 330.92 | 992.96 | 15956 |
| | Lentivirus | 107.78 | 81.70 | 88.31 | 51.84 | 133.73 | 12548 |
| | ACTB SLEEK | 388.65 | 365.80 | 166.62 | 284.86 | 456.85 | 10962 |

**Table S1. Level and variability of Optopatch expression across expression strategies.** FACS measurements in DIV28 NGN2 neurons of JF608 fluorescence and GFP fluorescence for PiggyBac-, lentivirus-, and ACTB-SLEEK-based Optopatch expression. Q1, 25th percentile; Q3, 75th percentile; N, number of analyzed cells. Related to Fig. S1.

| | Group | Mean | Median | Std | Q1 | Q3 | N |
|---|---|---|---|---|---|---|---|
| SNR | PiggyBac | 11.01 | 10.35 | 2.59 | 9.35 | 11.81 | 362 |
| | Lentivirus | 16.63 | 15.79 | 3.55 | 14.00 | 18.42 | 2053 |
| | ACTB SLEEK | 14.67 | 13.92 | 2.52 | 12.90 | 15.56 | 1046 |
| Total spikes | PiggyBac | 19.22 | 19.00 | 9.00 | 12.00 | 26.00 | 362 |
| | Lentivirus | 36.02 | 35.00 | 18.72 | 22.00 | 48.00 | 2053 |
| | ACTB SLEEK | 34.03 | 35.00 | 9.01 | 30.00 | 40.00 | 1046 |
| Optical rheobase | PiggyBac | 43.17% | 41.50% | 9.45% | 36.81% | 46.67% | 339 |
| | Lentivirus | 31.45% | 28.63% | 10.97% | 25.13% | 33.50% | 1875 |
| | ACTB SLEEK | 31.98% | 31.06% | 5.76% | 29.38% | 33.39% | 1033 |
| First spike rate (Hz) | PiggyBac | 5.73 | 5.52 | 1.72 | 4.57 | 6.67 | 299 |
| | Lentivirus | 8.96 | 7.84 | 11.78 | 6.35 | 9.30 | 1714 |
| | ACTB SLEEK | 8.09 | 6.90 | 10.61 | 6.15 | 7.96 | 996 |
| Last spike rate (Hz) | PiggyBac | 6.60 | 6.61 | 2.40 | 4.71 | 8.16 | 299 |
| | Lentivirus | 12.65 | 11.43 | 12.03 | 8.42 | 14.81 | 1714 |
| | ACTB SLEEK | 11.93 | 10.81 | 9.99 | 9.20 | 12.70 | 996 |
| Δspike rate (Hz) | PiggyBac | 0.87 | 1.23 | 2.34 | -0.36 | 2.31 | 299 |
| | Lentivirus | 3.69 | 3.69 | 16.81 | 1.40 | 6.39 | 1714 |
| | ACTB SLEEK | 3.85 | 3.89 | 14.55 | 2.33 | 5.34 | 996 |

**Table S2. Electrophysiological features across Optopatch expression strategies**. Electrophysiological features in DIV28 NGN2 neurons were compared among PiggyBac-, lentivirus-, and ACTB-SLEEK-based Optopatch expression. Metrics include spike signal-to-noise ratio (SNR), total number of detected spikes, optical rheobase (percent of maximum power to evoke the first spike during a blue-light ramp), initial and final spike rates during a blue-light ramp, and the change in spike rate during the ramp stimulus (Δspike rate). Q1, 25th percentile; Q3, 75th percentile; N, number of analyzed cells. Related to Fig. S3.

**Supplementary Movies**

**Movie 1:** Wide-area Optopatch recording of NGN2 neurons. Fluorescence fluctuations in each neuron are represented by the brightness of a randomly assigned color spot. Movie resolution 1,400 x 1,400 pixels, corresponding to 3.96 x 3.96 mm. Frame rate 800 Hz.

**Movie 2:** Wide-area Optopatch recording of NGN2 neurons. Fluorescence fluctuations in each neuron are represented by the brightness of a randomly assigned color spot. Movie resolution 2,000 x 2,504 pixels, corresponding to 5.65 x 7.08 mm. Frame rate 400 Hz.

**Movie 3:** Wide-area Optopatch recording of hiPSC-derived cardiomyocytes. Here the fluorescence fluctuations have been contrast-inverted so that depolarization appears as an increase in brightness. The cells were optogenetically paced at 1 Hz. Frame rate 500 Hz.

**Movie 4:** Wide-area Optopatch recording of hiPSC-derived cardiomyocytes.  Here the fluorescence fluctuations have been contrast-inverted so that depolarization appears as an increase in brightness.  The cells were optogenetically paced at 2 Hz. Frame rate 500 Hz.
